# Heterogeneity and plasticity of the naïve CD4^+^ T cell compartment

**DOI:** 10.1101/2024.10.02.615602

**Authors:** Alia Sajani, Evelien Schaafsma, Walburga Croteau, Mohamed ElTanbouly, Elizabeth C. Nowak, Chao Cheng, Christopher M. Burns, Mary Jo Turk, Randolph J. Noelle, J. Louise Lines

## Abstract

While naïve CD4^+^ T cells have historically been considered a homogenous population, recent studies have provided evidence that functional heterogeneity exists within this population. Using single cell RNA sequencing (scRNAseq), we identify five transcriptionally distinct naïve CD4^+^ T cell subsets that emerge within the single positive stage in the thymus: a quiescence cluster (T^Q^), a memory-like cluster (T^MEM^), a TCR reactive cluster (T^TCR^), an IFN responsive cluster (T^IFN^), and an undifferentiated cluster (T^UND^). Elevated expression of transcription factors KLF2, Mx1, and Nur77 within the T^Q^, T^IFN^, and T^MEM^ clusters, respectively, allowed enrichment of these subsets for further analyses. Functional studies using sorted cells revealed that naïve T cell subsets have distinctive functional biases upon stimulation. Furthermore, treatment of mice with inflammatory stimuli imparted a state of reduced responsiveness on naïve T cells, evidenced by a reduction in cytokine production *ex vivo*. In human lupus patients, naïve CD4^+^ T cell cluster frequencies were distorted, with the T^IFN^ cluster expanding proportionately with disease score. Our data show that naïve T cells are influenced by host environment, with functional consequences manifesting upon activation. These findings highlight a need to explore how naïve T cells can become distorted in cancer, autoimmunity, and infectious diseases.

**Summary:** This study describes the transcriptional heterogeneity of murine and human naïve CD4^+^ T cells as comprising of multiple discrete clusters that impact CD4^+^ T cell fate and trajectories. Naïve CD4^+^ T cells experiencing inflammatory environments exhibit an altered transcriptional state that influences their functional trajectory.

## Introduction

The naïve T cell state has been historically considered a homogenous, resting population (Van Den Broek et al., 2018). Naïve CD4^+^ T cells emerge from the thymus after undergoing negative and positive selection, enter the periphery, and circulate in a quiescent state before encountering antigen (Piconese et al., 2020; Vrisekoop et al., 2008). While circulating between the blood, spleen and lymph nodes – every 12 to 24 hours – naïve T cells encounter multiple environmental contexts (Jenni Punt VMT, 2013). A naïve T cell graduates to an effector T cell state after encountering its cognate antigen presented on an MHC molecule, along with engagement of costimulatory receptors on antigen presenting cells (APCs). The impact of the immune environment on the naïve T cell state has not been comprehensively characterized.

More recently, it has been shown that a number of well-defined signals control naïve T cell survival and fate, contributing to heterogeneity within the naïve T cell compartment. T cell survival is maintained by tonic TCR signaling (Surh and Sprent, 2008). Multiple studies have demonstrated how naïve T cells with varying degrees of tonic signaling exhibit functional heterogeneity when stimulated under identical conditions. Using Nur77-GFP reporting or antibody-based sorting of CD5^hi^ and Ly6C^lo^ to isolate high versus low tonic signaling naïve CD4s, one study showed that naïve CD4^+^ T cells that experience high tonic signaling are hyporesponsive and differentiate into Tregs, while two other studies demonstrated a Th17 lineage bias and greater IFNγ production in high tonically signaled CD4s (Martin et al., 2013; Henderson et al., 2015; Sood et al., 2019). In yet another enrichment of CD5^HI^ naïve T cells, the authors showed heightened propensity of these cells to differentiate into Tfh cells (Rogers et al., 2021). Notably, high tonic signaling also alters CD4^+^ T cell metabolism, which has the potential to affect the naïve T cell transition to an effector state (Milam et al., 2020). Therefore, tonic TCR signaling sustains naïve T cell homeostasis through not only maintaining naïve T cell survival, but also inducing a subset of naïve T cells that is highly sensitive to tonic TCR signaling, in turn influencing T cell function and differentiation.

Tonic IFN signaling sensitivity or responsiveness also impacts the naïve T cell state. Two studies isolated interferon signaling sensitive or responsive populations. Using the Tbet-reporter, a IFN sensitive master regulator for Th1 skewing, the authors of the study demonstrated that enriched naïve T cells acquired a heightened Th1 phenotype upon activation compared to their negative counterparts. When resting T cells were isolated using antibody-based sorting on BST2, an IFN-induced membrane protein, the BST2^HI^ CD4^+^ T cells produced more cytokines and had a heightened proliferative response to stimuli. In a published study on tonic IFN signaling in naïve CD8^+^ T cells, subsets of CD5^HI^/Ly6C^HI^ T cells exhibited a propensity to differentiate into CD8^+^ effector cells (Ju et al., 2021). Both IFN and TGFβ in naïve CD8^+^ T cells have been shown to alter CD8^+^ T cell functional responses post activation (Lee et al., 2023). Thus, tonic IFN signaling influences a subset of naïve T cells and alters their later functional trajectories.

Recent advancements in sequencing technologies have allowed for transcriptomic analyses of naïve CD4^+^ T cells. The consensus that all naïve cells comprise a functionally monomorphic population is challenged by our recent study, in which a redefinition of the naïve T cell state describes these cells as transcriptionally heterogeneous (ElTanbouly et al., 2020). Using single cell sequencing, we previously demonstrated an unexpected loss of quiescence in naïve CD4^+^ T cells from immune checkpoint VISTA (V-domain Ig suppressor of T cell activation) deficient mice compared to naïve CD4^+^ T cells with intact VISTA. The quiescence module was defined by *Klf2*, *Klf6*, *Btg1*, *Btg2* expression. Conversely, there was an expansion of the stem-cell memory-like cluster, defined by *Tcf7*, *Bcl2*, and *Il7r* (ElTanbouly et al., 2020). In addition to the finding that the immune checkpoint VISTA influences naïve T cells, as determined using single cell RNA sequencing, recent studies using such technologies also illuminated the role of cytokines in modulating the naïve T cell state. A published study showed that cytokines alone can polarize multiple immune cells, including T cells, in the lymph nodes of mice (Cui et al., 2024). Another study showed that elevated IL-4 presence during helminth infections results in the expansion of an IL4-induced naïve CD4^+^ T cell cluster, and these naïve CD4^+^ T cells are biased towards a hyporesponsive state upon alum treatment (Even et al., 2024). These findings set the stage for further characterization of heterogeneity within the naïve CD4^+^ T cell compartment.

This study explores how the immune environment shapes naïve CD4^+^ T cell transcriptional heterogeneity, and how these varied naïve T cell transcriptional states bias the differentiation trajectories of T cells once activated. Here, we propose a redefinition of the naïve T cell state to a compartment that is heterogenous; the naïve T cell program changes in different immune and disease contexts. Consequently, the steady-state heterogeneity of the naïve compartment has the potential to influence T cell activation and function, which could influence subsequent disease outcomes. Here, we identified 5 subsets in healthy adult mice using single cell sequencing of naïve CD4^+^ T cells and define how immune context impacts these naïve T cell clusters. These include a quiescence cluster (T^Q^), memory-like cluster (T^MEM^), TCR reactive cluster (T^TCR^), IFN responsive cluster (T^IFN^) and an undifferentiated cluster (T^UND^). This heterogeneity is imprinted in the thymus, maintained in the periphery, with the immune environment dynamically shaping T cell heterogeneity at the transcriptional level. We investigate how naïve CD4^+^ T cells maintain homeostasis during health and disease. While focusing on tonic TCR and IFN signaling, we explore immune contexts such as tissue, microbiome exposure and inflammation – and the influence these states have on naïve CD4^+^ T cells transcriptional identities. We also observe conservation of this transcriptional heterogeneity in human and mouse naïve CD4^+^ T cells, and demonstrate biased functional Th lineage commitment dependent on progenitor cluster identity. Further, we provide evidence supporting a redefinition of the T helper cell paradigm that renders the previously considered homogenous naïve T compartment, Th0, as heterogenous. This work highlights the importance for future studies to consider how heterogeneity of the naïve T cell compartment shapes susceptibility of the host to immune-related diseases such as infections, autoimmunity and cancer.

## Results

### Naïve CD4^+^ T cells are transcriptionally and functionally heterogeneous

Our prior study described changes in naïve CD4^+^ T cell heterogeneity caused by deficiency of the negative checkpoint protein VISTA (ElTanbouly et al., 2020). This study herein explores other mediators that influence naïve T cell heterogeneity, and how these differences within the naïve compartment modify T cell function and fate. To characterize naïve CD4^+^ T cell transcriptional heterogeneity, naïve CD4^+^ T cells (CD4^+^, CD8^-^, CD44^low^, CD62L^+^, CD25^-^, and CXCR3^-^) were bead enriched and FACS sorted to greater than 95% purity from spleens of C57BL/6J (WT) mice, and single cell sequencing (scRNAseq) was performed (Fig. 1A). In order to standardize results with our previous work, we clustered these data with our previously published VISTA-deficient naïve CD4^+^ T cells, which have a skewed naïve transcriptional profile that allows for optimal computational resolution of clusters (ElTanbouly et al., 2020). As shown by the UMAP in Figure 1B and frequency plot in Figure 1C, the naïve CD4^+^ T cell compartment was comprised of 5 transcriptionally-distinct clusters, distinguished by marker genes shown in Figure 1D. The T^Q^ (quiescence) cluster expressed genes associated with T cell quiescence (*Klf2*, *Klf6*, *Jun* and *Btg2)* (Fig. 1D; Fig. S2). T^MEM^ (memory-like) cluster expressed higher levels of genes that are generally associated with memory stemness (*Ccr7*, *Il7r* and *Tcf7)* (Fig. 1D; Fig. S2). In contrast, the T^TCR^ cluster was marked by genes induced by TCR signaling such as *Cd5*, *Cd6* and *Nr4a1* (Fig. 1D; Fig. S2), potentially signifying recently experienced tonic TCR signaling or higher levels of baseline tonic signaling. The cluster termed T^IFN^ was defined by expression of Interferon-Stimulated Genes (ISGs) including *Stat1*, *Irf7* and *Ifit1* (Fig. 1D; Fig. S2), whereas a final cluster expressed few unique marker genes, but displayed a modest enrichment of cytoskeletal genes (e.g. *Act1g)*, and was therefore termed the T^UND^ (undifferentiated) (Fig. 1D; Fig. S2). In general, the T^Q^, T^MEM^, and T^IFN^ clusters appeared closely related to those found in our prior study, with the T^UND^ cluster being similar to our previously reported extracellular matrix and cytoskeletal-like cluster (ElTanbouly et al., 2020). These data highlighted the presence of heterogeneity within the naïve CD4^+^ T cell compartment, although it remained unknown whether these clusters also had divergent fate or functional diversity after activation.

**Figure 1.**
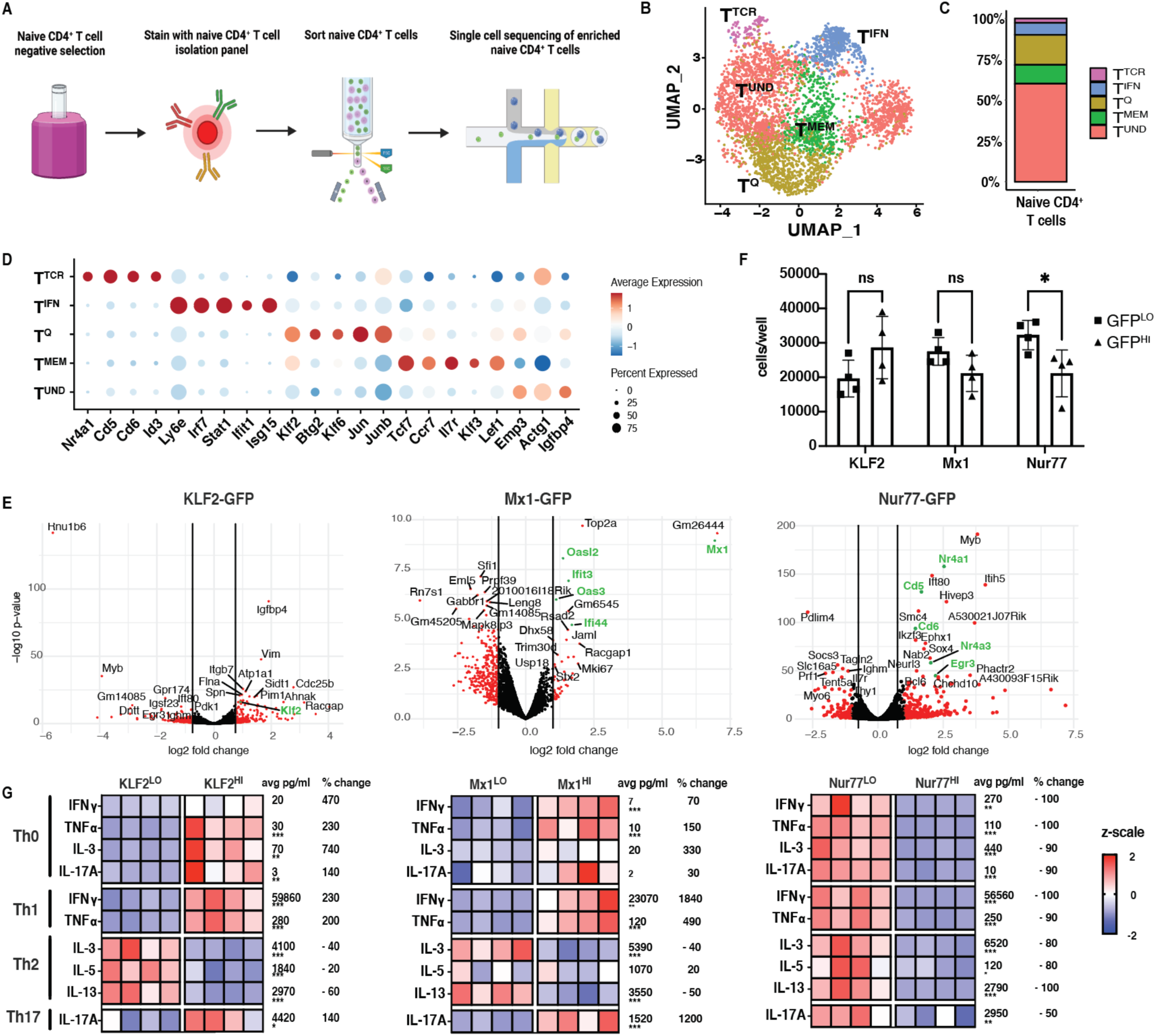
Naïve CD4^+^ T cells are transcriptionally and functionally heterogeneous. **A:** Experimental outline for enrichment of naïve CD4^+^ T cells. **B:** UMAP of naïve CD4^+^ T cells from C57BL/6J spleens colored by cell cluster based on Louvain clustering (n=2). **C:** Stacked bar plot of frequencies in (B), grouped by cluster. **D:** Dot plot of cluster-defining marker genes for cell type annotations. Dot size represents fraction of cells expressing a gene in each cluster. Dot color represents scaled average expression by gene column. Red depicts gene upregulation while blue represents gene downregulation. **E:** Volcano plots showing bulk sequencing of sorted of naïve CD4^+^ T cells sorted from top and bottom 10-15% GFP reporter mice, denoted as Nur77^HI^ or Nur77^LO^ and KLF2^HI^ or KLF2^LO^. For Mx1^HI^, naïve CD4^+^ T cells with positive GFP expression (top 1%) was collected, while the bottom 10-15% was collected for Mx1^LO^. The x axis is the log2fold change value and the y axis is −log10 of the p value. Genes labeled in green correspond to cluster markers, with at least 2 samples per group. **F:** Proliferation of Nur77^HI^ vs Nur77^LO^, KLF2^HI^ vs KLF2^LO^ and Mx1^HI^ vs Mx1^LO^ naïve CD4^+^ T cells after 5 days of anti-CD3/28 activation and culture in Th1 conditions (n=4). **G:** Heat maps showing cytokine production in pg/ml from Luminex data from sorted Nur77^HI^ vs Nur77^LO^, KLF2^HI^ vs KLF2^LO^ and Mx1^HI^ vs Mx1^LO^ naïve CD4^+^ T cells activated and cultured in Th0, Th1, Th2 and Th17 skewing conditions for 5 days. Scale is displayed as z scale normalization, with average numbers across all groups displayed to the right of each row (n=4). Data representative of at least 2 experiments per reporter strain. Statistical significance testing by multiple unpaired t-tests. * <0.05, ** <0.01, ***<0.001. Percent change denotes change in cytokine production from reporter low group to reporter high group.

To address if functional distinctions exist amongst resting T cell subsets defined by scRNAseq, efforts were implemented to enrich these subsets for functional studies. CITEseq was used in an attempt to identify cell surface markers for naïve CD4^+^ T cells that might define single cell transcriptional clusters of resting T cells. Naïve CD4^+^ T cells isolated from murine spleen or human peripheral blood were stained with a cocktail of 119 (mouse) or 130 (human) CITEseq mAbs and were then clustered based on mRNA expression. Supplemental Figure S3 shows CITE-Seq data from murine naïve CD4^+^ T cells, while Supplemental Figure S4 shows CITE-Seq data from human naïve CD4^+^ T cells. The murine T^TCR^ cluster expressed a modest increase in surface protein expression of FR4, CD5 and CD69 (Fig. S3). However, no additional surface phenotypes emerged to allow for potential flow cytofluorimetic sorting and enrichment of other scRNAseq defined clusters (Fig. S3-4). In some cases, protein and RNA expression did not correlate (Fig. S3). For example, for CD127, transcriptional expression was elevated in the T^MEM^ cluster, but protein was enriched in other clusters (Fig. S3). Predicted protein markers of interest from human naïve CD4^+^ T cell CITE-Seq data also failed to overlap with transcriptional signatures (Fig. S4). The antibodies in the CITEseq cocktail thus did not reveal surface proteins that could be used to sort-purify clusters.

However, the clusters identified by scRNAseq were noted for their heightened expression of specific transcription factors (TF). Therefore, an alternative strategy was used in which naïve CD4^+^ T cells were sorted from TF reporter mouse strains based on high or low expression of reporter protein. Mice expressing Nur77-GFP were used to enrich for naïve T cells with a T^TCR^ signature, based on their high expression of *Nr4a1*. Naïve T cells expressing high levels of GFP (highest 15% MFI GFP; designated Nur77^HI^) or low levels of GFP (lowest 15% MFI GFP; designated Nur77^LO^) were sorted (Fig. S5), allowing for sufficient numbers of cells to be functionally evaluated. T^Q^, previously characterized as a highly quiescent naïve CD4^+^ T cell subset, was marked by *Klf2*. Thus, KLF2-GFP reporter mice were used in an attempt to enrich the T^Q^ cluster, using a similar reporter sorting approach of top and bottom 15% MFI GFP expression (Fig. S5). Finally, Mx1-GFP reporter mice, which report type I and type III interferon responsiveness, were used to enrich for T^IFN^, by sorting on GFP^+^ (top 1%) of naïve CD4s (designated Mx1^HI^), with the lowest 15% MFI GFP designated as Mx1^LO^ (Fig. S5).

To validate that GFP reporter sorting effectively enriched for our transcriptionally-defined clusters, sorted cell populations were analyzed by bulk RNA seq. Indeed, Nur77^HI^ naïve T cells demonstrated enriched expression of T^TCR^ cluster genes (e.g. *Nr4a1, CD5* and *CD6*) while Nur77^LO^ cells showed diminished expression of T^TCR^ cluster genes (Fig. 1E). Similarly, Mx1^HI^ naïve CD4^+^ T cells expressed higher levels of T^IFN^ markers *Mx1*, *Ifi44, Ifit3, Oas3* and *Oasl2*, and KLF2^HI^ cells expressed higher *Klf2* (Fig. 1E). Of note, while we had also attempted to use TCF7-GFP mice to isolate the T^MEM^ population, but found that genes in that cluster were not enriched in TCF7^HI^ sorted cells (Figure. S7). Thus, using TF reporter mice, we were able to sort populations that represented three of the major subsets within the naïve CD4^+^ T cell compartment.

To functionally assess differences in naïve differentiation capabilities, after naïve CD4^+^ T cell subsets were enriched based on TF-GFP expression, their functional capacities were evaluated in vitro. First, we examined proliferative capacities upon anti-CD3/CD28 activation, under Th1 skewing conditions. While proliferation of Mx1^HI^ vs Mx1^LO^, and KLF2^HI^ vs KLF2^LO^ naïve T cell subsets was similar, Nur77^LO^ cell proliferation significantly exceeded that of Nur77^HI^ cells (Fig. 1F). The reduced proliferation of the T^TCR^ cluster is supported by a prior study that reported that highly self-reactive naïve CD4^+^ T cells are biased towards a Treg phenotype (Martin et al., 2013).

To further identify skewing trajectories of each population, T cells were activated (with anti-CD3/anti-CD28) for 5 days in either Th0, Th1, Th2, or Th17 skewing conditions, and cytokines production was quantified by Luminex. Comparing KLF2^HI^ and KLF2^LO^ naïve CD4^+^ T cells under Th1-skewing conditions, IFNγ production differed by a factor of 3.3-fold (from an average of 28,000 vs. 92,000 pg/ml), and TNFa production was increased by 2-fold (Fig. 1G). A similar but less pronounced trend was observed in Th17 culturing conditions, where KLF2^HI^ cell average IL-17A production was increased by 1.4-fold (2622 vs 6220 pg/ml) relative to their KLF2^LO^ counterparts (Fig. 1G). Conversely, in Th2 conditions, KLF2-GFP^HI^ cells made between 20-60% less IL-3 (5043 vs. 3159 pg/ml), IL-5 (2100 vs. 1600 pg/ml) and IL-13 (4300 vs. 1700 pg/ml) compared with KLF2^LO^ cells (Fig. 1G). These data support the conclusion that KLF2^HI^ T^Q^ cells preferentially adopt a Th1 or Th17 fate, but resist adoption of the Th2 fate.

In assessing the skewing potential of Mx1^HI^ cells, a similar pattern was observed. IFNγ production increased 1840% and TNFa production increased 490% in naïve Mx1^HI^ cultured in Th1 skewing conditions compared to Th1 cultured naïve Mx1^LO^ cells (Fig. 1G). In Th17 conditions, IL-17A production by Mx1^HI^ cells was 12-fold higher (219 vs 2831 pg/ml) (Fig. 1G). However, in Th2 conditions, average IL-3 production by Mx1^HI^ cells was reduced by ∼40% (6600pg/ml vs. 4200 pg/ml) and IL-13 production was ∼50% lower (4800 vs. 2300 pg/ml) (Fig. 1G). Thus, similar to T^Q^ cells, Mx1^HI^ T^IFN^ cells skewed away from a Th2 fate, preferentially producing Th1 and Th17 cytokines.

Finally, activation and skewing of Nur77^HI^ naïve CD4^+^ T cells revealed reduced cytokine production of ∼50-100% across all skewing groups – including IFNγ, TNFα, IL-3, IL-13 and IL-17a – when compared to Nur77^LO^ naïve CD4^+^ T cells (Fig. 1G). In Th1 conditions, IFNγ production decreased in Nur77^HI^ naïve CD4^+^ T cells (110000 pg/ml to 1800 pg/ml). For Th2 culturing conditions, there was also a decrease in IL-3 (11000 to 1800 pg/ml) and IL-5 (200 to 43 pg/ml) (Fig. 1F). When the Nur77^HI^ naïve CD4^+^ T cells were cultured in Th17 conditions, IL-17A production dropped (4010 pg/ml to 1900 pg/ml), compared to Nur77^LO^ naïve CD4^+^ T cells (Fig. 1G). Taken together with the low proliferative capabilities of Nur77-GFP^HI^ naïve CD4^+^ T cells (Fig. 1E), these data are consistent with the idea that T^TCR^ cells have elevated tonic signaling and anergy marker expression (such as FR4, as shown by CITE-Seq in Fig. S3), and suggest that the T^TCR^ cluster is a hyporesponsive subset of naïve CD4^+^ T cells, potentially with an immunoregulatory program that prevents self-reactivity. These studies establish a predetermined and varied fate of naïve CD4^+^ T cell subsets which, despite being uniformly activated, display functional heterogeneity.

### Naïve CD4^+^ T cell heterogeneity is established in the thymus

To characterize the ontogeny of the naïve CD4^+^ T cell clusters, their representation within thymic T cell subsets was investigated. The 5 transcriptional clusters of resting CD4^+^ T cells were incompletely represented in the CD4^+^ CD8^+^ double positive (DP) lineage, but all emerged in the CD4^+^ CD8^-^ single positive (SP) lineage (Fig. 2A). These clusters were maintained in recent thymic emigrants (RTEs) and peripheral naïve CD4s (Fig. S9 and Fig. 2A). At the double positive (DP) stage, the T^TCR^ frequency was 16% and T^Q^ frequency was 5% (Fig. 2A and Fig. 2B). At the single positive (SP) stage, the frequency of T^TCR^ increased to 39% and T^Q^ increased to 21% (Fig. 2A and Fig. 2B). T^MEM^ frequencies also increased from 3% to 25% in the DP to SP transition, while T^IFN^ increased from 2% to 55%, illustrating an emergence of these clusters during the DP to SP transition (Fig. 2A and Fig. 2B). mTECs (medullary Thymic Epithelial Cells) are the main source of type I and III IFNs in the thymus, and may be the driving force imprinting T^IFN^ in the thymus (Benhammadi et al., 2020; Martinez et al., 2023). Violin plots show emergence of *Cd5, Nur77, Stat1, Klf2, Klf6, Il7R* and *Ccr7* in the SP thymocyte development stage (Fig. 2C). As expected for cells early in T cell development, at the DP stage, the majority of cells expressed developmental genes such as *Rag1, Rag2, Dntt, Myb* and *CD24a* (Fig. S9), referred to herein as the Immature clusters (Kernfeld et al., 2018). The DP T^UND^ cells did not completely overlap with SP T^UND^ or splenic T^UND^ from CD4^+^ T cells in UMAP space, suggesting that the T^UND^ in DP and SP were distinctive (Fig. 2A). The thymic environment, as well as the T cell developmental transition from DP to SP, was critical to the establishment of the naïve CD4^+^ T cell clusters. The thymus is therefore a critical site of naïve CD4^+^ T cell development that extends beyond the classically defined negative and positive selection mechanisms; controlled developmental cues within the thymus impart transcriptional and functional diversity through the establishment of distinct naïve T cell subsets.

**Figure 2.**
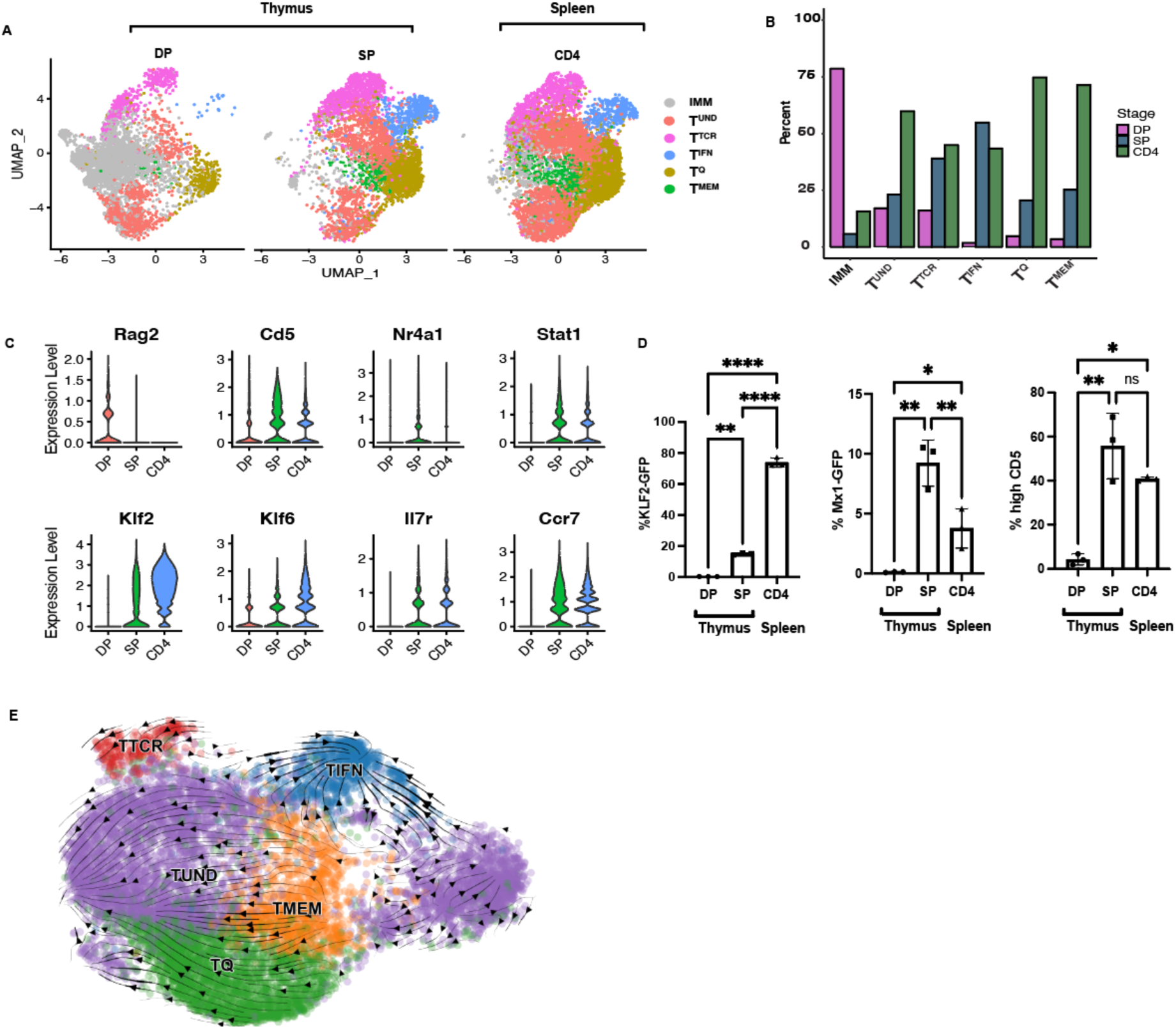
Ontogeny of resting T cell clusters. **A:** UMAP of SP CD4^+^ and DP CD4^+^CD8^+^ thymocytes and naïve CD4^+^ T cells in the spleen. Mice used were VISTA CD4-Cre-, representative of wild type mice. (DP: n=1, SP: n=2, CD4: n=3). **B:** Stacked bar plot of frequencies in (A), grouped by cluster. **C:** Violin plot of marker genes (A). **D:** Mx1-GFP and KLF2-GFP reporting, and CD5 expression, in SP and DP thymocytes, and peripheral naïve CD4^+^ T cells. Significance testing was performed using two-way ANOVA. * <0.05, ** <0.01, ***<0.001. **E:** RNA velocity analysis of splenic naïve CD4^+^ T cells from 2 C57BL/6J donors, plotted on the same UMAP as in Fig. 1. Arrows represent potential differentiation paths calculated using velocity vectors from unspliced to spliced RNA ratios.

To validate the emergence of the 5 clusters in the thymus, we used reporter strains and surface marker staining to identify our clusters. In Mx1-GFP mice, GFP was not present in DP, but was expressed in 9% of naïve CD4s in the SP stage and in 4% of peripheral CD4s (Fig. 2D), validating the emergence of T^IFN^ during the SP to DP transition. CD5 is a marker of TCR reactivity and is expressed on all T cells, with the intensity of CD5 staining higher the T^TCR^ cluster (Azzam et al., 1998). This data as well as our CITE-Seq data (Fig. S3-4) enabled us to use CD5^hi^ expression as a marker gene for the T^TCR^ cluster. High CD5 was observed on 4% of DP cells, but on 56% of SP cells, which have completed positive selection (Fig. 2D). Similarly, T^Q^, represented by KLF2-GFP, also confirmed the emergence of T^Q^ in the DP stage, with significantly increased GFP reporting in peripheral CD4s (Fig. 2D). Thus, as thymocytes reached the SP stage of selection, naïve CD4 heterogeneity was comprehensively imprinted, and this transcriptional program was maintained in the periphery (Fig. 2A-B). Thymic education therefore guides naïve CD4^+^ T cell subset formation, with each subset adopting varied gene signatures.

As a computational approach to exploring the ontogeny of naïve CD4^+^ T cell clusters, we employed RNA velocity, which provides temporal information based on the ratio of spliced vs unspliced RNA. Fig. 2E shows the RNA velocity of naïve CD4^+^ T cells, with the arrows showing predicted directional development of, and relationships between, naïve CD4^+^ T cell transcriptional clusters. The short RNA velocity lines depict a relative stability of the clusters, as well as the low transcriptional activity of naïve CD4^+^ T cells. Despite the low gene expression pattern, we observed T^IFN^ as a terminally differentiated cluster. RNA velocity arrows converge into the T^IFN^ cluster, suggesting that T^IFN^ is a terminal cluster (Fig. 2E). Supplemental Figure S10 shows RNA velocity, PAGA (Partition-based graph abstraction) and pseudotime plots, further demonstrates the dynamic temporal relationship between the naïve CD4^+^ T cell clusters (Wolf et al., 2019). T^Q^ and T^UND^ emerge from T^MEM^, while T^TCR^ is a more stable cluster, depicted by the circulating arrows (Fig. 2E). Therefore, the RNA velocity analyses suggest that the naïve CD4^+^ T cell cluster frequencies may be homeostatically controlled, yet could have the potential to be plastic, which would result in some clusters giving rise to other more differentiated naïve CD4^+^ T cell subsets. This transcriptional homeostasis between stability and plasticity of CD4^+^ T cell gene expression may allow for the survival and maintenance of the naïve cell state.

### Impact of TCR signaling on naïve CD4^+^ T cell heterogeneity

TCR affinity is known to impact on T cell identity and functional diversity (This et al., 2021; Martinez and Evavold, 2015). To determine the influence of TCR specificity on imprinting naïve CD4^+^ T cell heterogeneity, we evaluated transcriptional clusters in resting T cells from OTII and SMARTA TCR transgenic mouse strains (Fig. S11). The scRNAseq determined distribution of naïve clusters in both TCR Tg strains was similar to the C57BL/6J control (Fig. S11). However, SMARTA mice, a strain that has been previously reported to have heightened CD5 expression as well as more self-reactive T cells (Mandl et al., 2013), had a slightly larger T^TCR^ cluster (Fig. S11). Regardless, the frequency of T^TCR^ cells was not significantly different between polyclonal (C57BL/6J) and monoclonal (OTII and SMARTA) strains, or between the monoclonal strains. Thus, antigen specificity does not prominently influence cluster heterogeneity.

Maintenance of the peripheral resting T cell compartment is dependent on tonic TCR signaling. As such, we hypothesized that tonic signaling may also impact the steady state heterogeneity of resting T cells in the periphery. To address this hypothesis, tonic signaling was blocked in vivo using anti-MHC II administration (1mg/mouse every day for 3 days, with the takedown on Day 4, as previously reported (Stefanová et al., 2002), and the heterogeneity of naïve CD4^+^ T cell clusters was evaluated. This short-term treatment with anti-MHC II was not predicted to globally impact on T cell numbers, as naïve CD4^+^ T cell numbers do not diminish over one month in MHC Class II deficient hosts (Dorfman et al., 2000). However, treatment with anti-MHC Class II resulted in a significant reduction in the frequency of the T^TCR^ cluster (3-5% to 0.4-0.8%) compared with isotype control (Fig. 3A-B). This reduction was unique to the T^TCR^, establishing the existence of T^TCR^ as uniquely dependent on tonic signaling. We also observed a global decrease in TCR signaling genes amongst all naïve T cells from the anti-MHC II treated mice compared to isotype control (Fig. 3C) which was anticipated as all T cells experience some tonic TCR engagement. These findings were validated using flow cytometry staining of naïve Nur77-GFP reporting CD4^+^ T cells, which represent T^TCR^, having lower Nur77-GFP production with anti-MHC II treatment (Fig. S12). The short period of treatment did not result in substantial T cell death, as there was no statistically significant difference in Annexin V staining between anti-MHC II treated and isotype treated mice (Fig. S12). Further, GSEA analysis showed that MAP2K and MAPK activating pathway genes were also globally reduced in anti-MHC II treated naïve CD4s compared to isotype controls, and CD28 costimulation-related genes were also reduced (Fig. 3D). Despite naïve CD4s not experiencing downstream costimulation receptor signaling, blocking MHC II-TCR interactions may prevent interactions between naïve T cells and APCs that are needed for tonic signaling of costimulatory pathways (Shah et al., 2021). GSEA analysis also showed an increase in interferon signaling pathways (Fig. 3D). It is possible that naïve CD4s in the T^TCR^ cluster lose their T^TCR^ signature and take on a T^IFN^ signature due to the increase in Mx1-GFP reporting in anti-MHC II and isotype treated mice (Fig. S12). Alternatively, cross-linking of MHCII with anti-MHC Class II treatment may have resulted in the observed increase of IFN genes. Thus, tonic signaling maintains naïve CD4^+^ T cell homeostasis to varying degrees across naïve T cell subsets, with the greatest influence on T^TCR^. The naïve CD4^+^ T cells in this cluster presumably have more self-reactive TCRs and are either generated by tonic signaling or highly sensitive to loss of tonic signaling compared to other naïve CD4^+^ T cell subsets.

**Figure 3.**
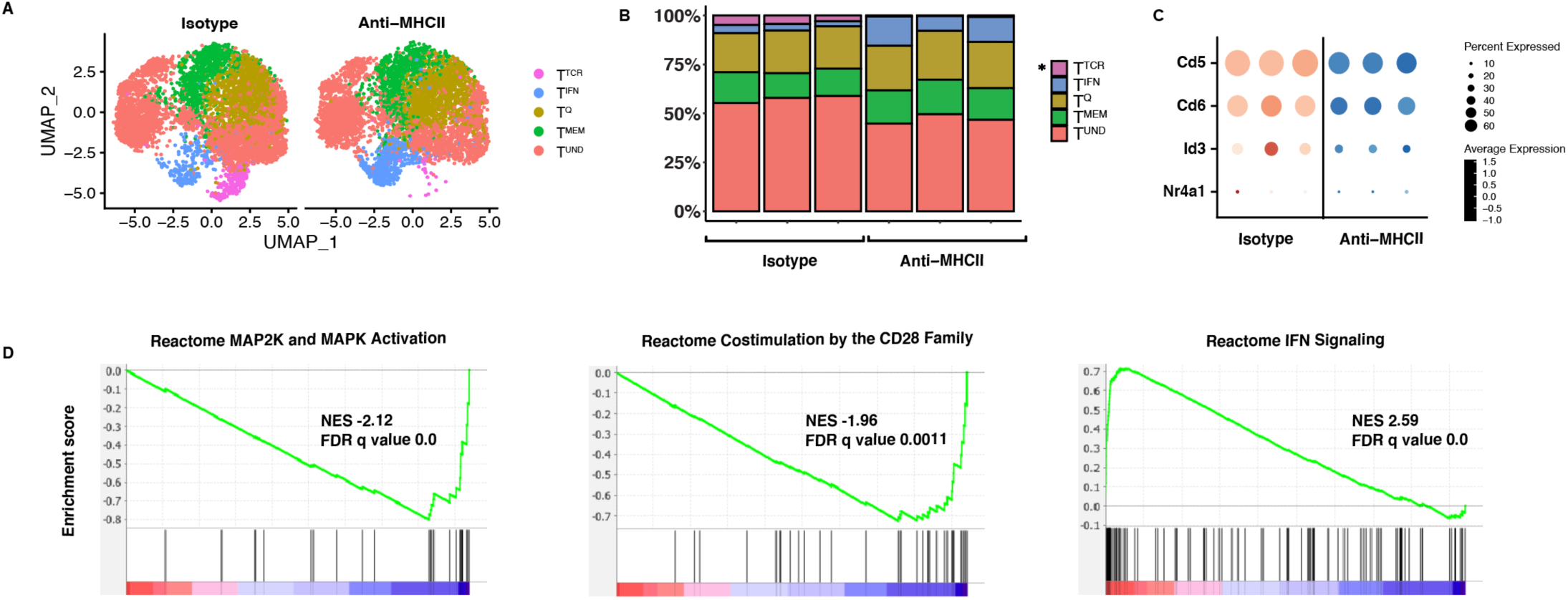
Tonic signaling impacts the T^TCR^ naïve CD4^+^ T cell transcriptional cluster. **A:** UMAP of naïve CD4^+^ T cells from C57BL/6J mice treated with anti-MHCII or isotype control (via i.p. injection for 3 consecutive days with 1mg/mouse, with the takedown on day 4). **B:** Stacked bar plot of frequencies in (A). **C:** Dot plot of T^TCR^ genes comparing naïve CD4^+^ T cells from anti-MHCII treated and isotype treated mice. **D:** GSEA analysis of anti-MHCII treated and isotype treated naïve CD4^+^ T cells, showing MAP2K/MAPK activation, costimulation by CD28 family and IFN signaling gene signatures anti-MHCII blockade. (n=3).

### Tissue microenvironment shapes naïve CD4^+^ T cell heterogeneity

Since naïve CD4^+^ T cells continually recirculate through the blood and secondary lymphoid organs (SLOs), naïve CD4^+^ T cells experience distinct tissue environments. Therefore, we assessed whether naïve CD4^+^ T cell transcriptional heterogeneity changes across various tissues. In C57BL/6J mice, we observed an increase in the size of the T^Q^ and T^MEM^, and decrease in T^TCR^, in blood compared with the spleen (Fig. 4A and 4B). To test the impact of tissue microenvironment experience on naïve CD4^+^ T cell gene expression, we adoptively transferred congenically-marked (CD45.1+) naïve CD4^+^ T cells isolated from blood – and isolated these CD45.1+ naïve CD4 T cells that made their way through the circulation and into the spleens of the recipient mice 3 days later (Fig. 4C). scRNAseq revealed that blood cells which localized to the spleen clustered with comparable proportions to endogenous naïve CD4^+^ T cells in spleens of recipient mice (Fig. 4D). Upon statistical testing, we observed a statistically significant difference between naïve CD4^+^ T cells from the blood and spleen, for both the T^Q^ and T^MEM^ clusters (p = 0.99 and 0.94 respectively), but no statistical difference between the blood transferred cells in the spleen and host splenic naïve CD4^+^ T cells. Transferred cells also lost the quiescence signature found in blood (Fig. 4E). The difference in T^TCR^ and T^IFN^ frequencies between spleen and blood were not statistically significant (Fig. 4D). Notably, the transferred CD45.1 blood naïve CD4^+^ T cells, acquired a migratory gene expression similar to splenic naïve CD4^+^ T cells (Fig. 4F). The transferred CD45.1 blood naïve CD4^+^ T cells downregulated *S1pr1, Ccr7* and *Lgals9* in the spleen (Fig. 4F). We also observed higher expression of *Rhoa, Itgb7* and *Coror1a*, in the transferred cells, even compared to host spleen, indicating a homeostatic shift that is later tempered when the naïve CD4^+^ T cells undergo prolonged retention in the spleen (Fig. 4F). Therefore, changes in transcriptional heterogeneity of naïve CD4^+^ T cells, due to differences in tissue microenvironment, highlights the plasticity of the naïve T cell compartment. However, an alternative explanation for the equivalence in cluster frequencies between host spleen and transferred blood in the spleen is selective survival or recruitment of naïve CD4^+^ T transcriptional subsets to the spleen.

**Figure 4.**
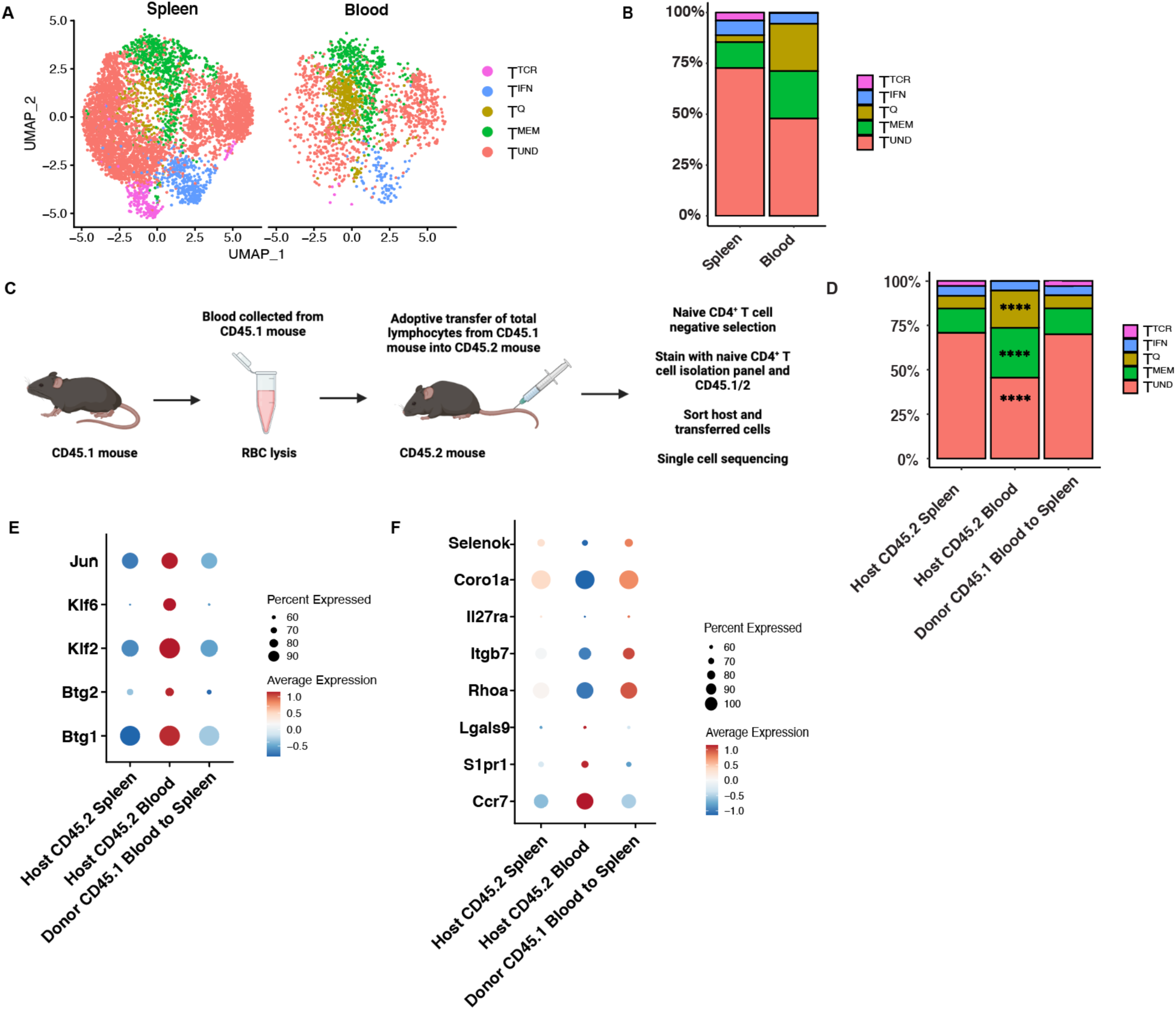
Tissue microenvironment dependent homeostasis of naïve CD4^+^ T cells in the spleen and blood. **A:** UMAP of naïve CD4^+^ T cells from C57BL/6J mice spleens, lymph nodes and blood. (n=2). **B:** Stacked bar plot of frequencies in (A). **C:** Experimental outline of adoptive transfer of lymphocytes from blood into host mouse, followed by isolation and sorting of transferred blood naïve CD4^+^ T cells from host and donor spleen and blood after 3 days. (n=2). **D:** Stacked bar plot of frequencies in from adoptive transfer experiment in (C). All statistical significance testing in figure was performed using 2-way ANOVA. * <0.05, ** <0.01, ***<0.001. **E:** Dot plot of T^Q^ genes in adoptive transfer experiment. **F:** Dot plot of migratory genes in adoptive transfer experiment.

### Naïve CD4^+^ T cells exhibit skewing bias after inflammation experience

A major tenant of immune responses is regulation of inflammation, which can include multiple types of inflammation. The microbiome has been shown to alter immunological responses in many studies (Akrami et al., 2020; Wiertsema et al., 2021). One study reported success of anti-PD1 immunotherapy for melanoma, in part due to altered T cell effector potentials, depends on microbial composition (Gopalakrishnan et al., 2018). Another study showed that CX3CR1^+^ dendritic cells transport microbial peptides from the intestines to the thymus for tolerance induction (Zegarra-Ruiz et al., 2021). We sought to determine whether naïve CD4^+^ T cells would have an altered transcriptional profile in germ free mice, due to the lack of microbial cues that could potentially shape the construction of the naïve T cell compartment. We performed single cell RNA sequencing on sorted naïve CD4^+^ T cells from germ free mice obtained from Charles River (C57BL/6-Germ-Free). In order to have the appropriate Wild Type control with the same genetic background, we also performed single cell sequencing on C57BL/6 mice from Charles River. We found no statistically significant difference in cluster frequencies between the germ free and specific pathogen free naïve CD4^+^ T cells (Fig. S13). The observed lack of microbiome dependent influence on naïve CD4^+^ T cell transcriptional heterogeneity may be due to the microbial composition and housing conditions of specific pathogen free mice, or may be because naïve CD4^+^ T cells have not yet encountered microbes.

Even though CD4^+^ T cells aren’t changed in the absence of an intact microbiome, potent inflammatory insults may impact the naïve CD4^+^ T cell state. Prior studies on the impact of sepsis on naïve CD4^+^ and CD8^+^ T cells demonstrated a reduction in the frequency of naïve T cells during potent inflammation (Markwart et al., 2014; Condotta et al., 2013). We designed a study to address whether the induction of an acute, inflammatory environment, created by LPS or CFA administration, alters the transcriptional profiles within the naïve CD4^+^ T cell compartment. LPS or CFA were administered to mice, and after 7 days, naïve CD4^+^ T cells were isolated and subjected to scRNAseq. Frequencies of the T^TCR^ and T^MEM^ naïve T clusters were not significantly different between naïve T cells from CFA or LPS vs untreated mice (Fig. 5A and 5B). However, T^IFN^ was significantly lower with both CFA (p = 0.0044) and LPS (p = 0.0005) treatment, as compared to untreated, and similar between the two treatment groups (Fig. 5B). T^Q^ was also significantly lower with CFA (p = 0.0040) but not with LPS treatment (p = 0.14) (Fig. 5B). Upon differential gene expression analyses across the clusters, we observed an inflammation-induced skewing of naïve CD4^+^ T cell transcriptional states (Fig. 5C). ISGs, TNFα/NFkB inhibitory genes and quiescence-associated genes were downregulated upon experience of a CFA inflammatory environment (Fig. 5C). Conversely, self-renewal/stemness-related, ribosomal and elongation genes were upregulated (Fig. 5C). The naïve CD4^+^ T cells on day 7 may be downregulating IFN responsiveness sustain the naïve T cell state, since peak IFNγ production has been reported to be between 6-8 hours (Varma et al., 2002). Therefore, inflammation altered bystander naïve CD4^+^ T cells at the transcriptional level.

**Figure 5.**
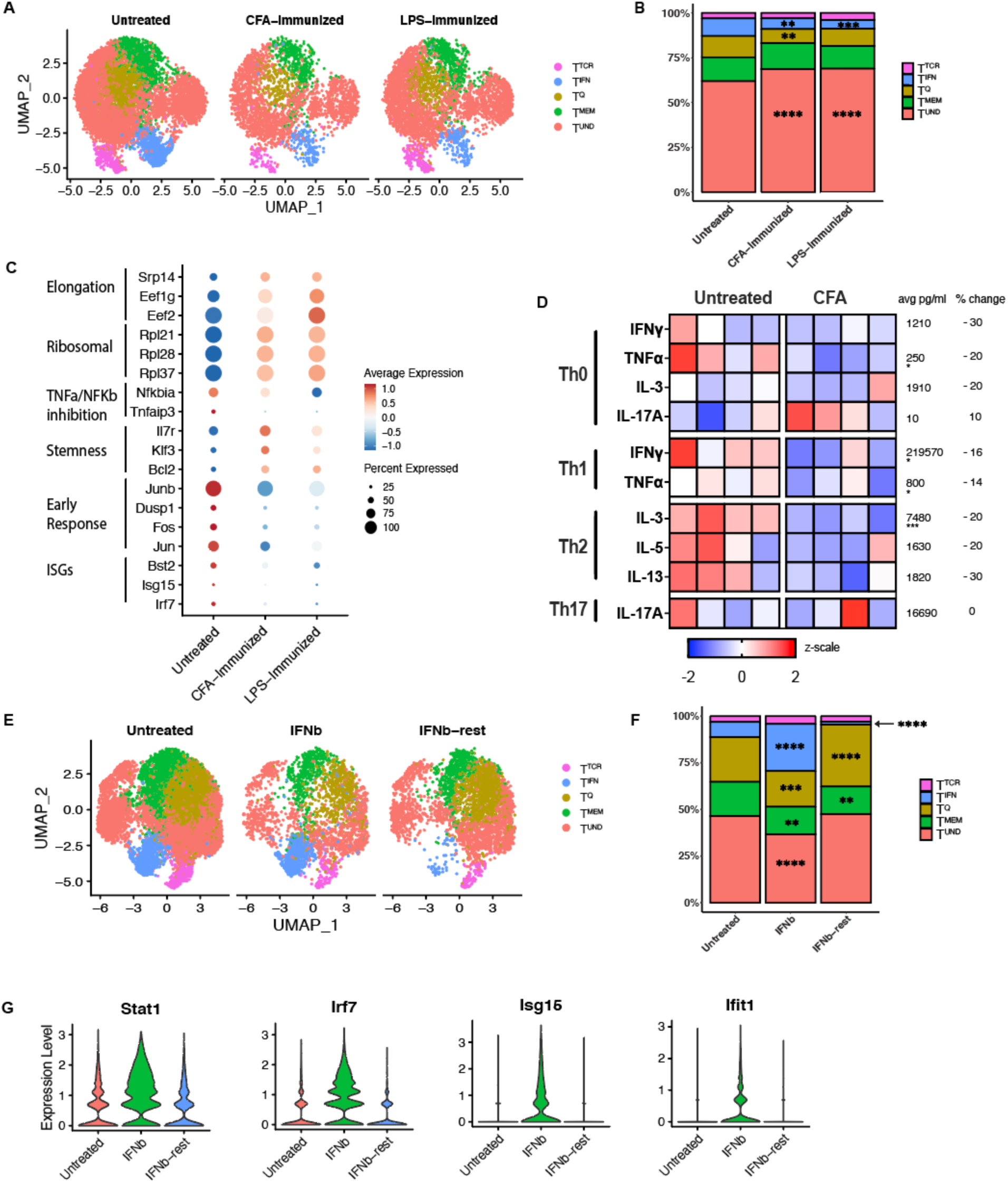
An inflammatory environment alters naïve CD4^+^ T cell transcriptional and functional states. **A:** UMAP of naïve CD4^+^ T cells from spleens isolated from CFA treated (injected i.p. with a 1:1 emulsion of 100ug CFA and PBS or 10ug LPS, takedown on day 7) C57BL/6J mice and untreated controls. (n=2). **B:** Stacked bar plot of frequencies in (A). All statistical significance testing in figure was performed using 2-way ANOVA. * <0.05, ** <0.01, ***<0.001. **C:** Differentially expressed genes across CFA treated, LPS treated and untreated control mice. **D:** Skewing trajectories of naïve CD4s from CFA treated (after 7 days) and untreated control mice that were activated and cultured in Th0, Th1, Th2 and Th17 skewing conditions for 5 days. Data representation of at least two experiments. Scale is displayed as z scale normalization, with average numbers across all groups displayed to the right of each row. Percent change denotes change in cytokine production from untreated group to CFA treated group. **E:** UMAP of naïve CD4^+^ T cells from untreated mice, mice treated with 0.25ug IFNβ daily for 5 days, or mice treated with 0.25ug IFNβ daily for 5 days followed by a 5 day of rest. (n=2). **F:** Stacked bar plot of frequencies in (E). All statistical significance testing in figure was performed using 2-way ANOVA. * <0.05, ** <0.01, ***<0.001. **G:** Violin plots for ISGs in untreated, IFNβ treated or IFNβ treated and rested naïve CD4^+^ T cells.

Next, we investigated if the administration of an inflammatory challenge elicited differences in resting T cell functional fates. When cultured in Th0, Th1, Th2 and Th17 conditions, naïve CD4^+^ T cells from CFA treated mice were less responsive compared to naïve CD4^+^ T cells from untreated mice. In Th1 culturing conditions, IFNγ production was reduced 30% from 240,000 pg/ml to 200,000 pg/ml, and TNFa production 20% from 860 pg/ml to 740 pg/ml, when comparing naïve T cells from untreated vs. CFA treated mice (Fig. 5D). In Th2 culturing conditions, IL-3 production was reduced 20% from 8400pg/ml to 6600pg/ml (Fig. 5D). Therefore, induction of an inflammatory environment generally reduced the responsiveness of naïve CD4^+^ T cells, consistent with the idea that acute inflammation restricts naive T cell activation.

Then, instead of exploring global inflammation induced by CFA, we sought to investigate the impact of a single inflammatory cytokine on heterogeneity. So, we treated mice with IFNβ and assessed their phenotype after 5 days of continuous treatment or after 5 days of treatment and 5 days of rest. IFNβ administration resulted in an increase in the frequency of cells defined as T^IFN^ cluster, expanding from 8% to 24-26%. However, the treated and rested naïve CD4^+^ T cells contained a T^IFN^ cluster than was significantly smaller even than that of untreated mice, at a frequency of 2% (Fig. 5F). Moreover, violin plots (Fig. 5G) demonstrate individual ISG increase in IFNβ treated mice, that were quickly reduced in IFNβ treated and rested mice. This expansion and contraction of the T^IFN^ cluster is consistent with published data showing that IFN response is followed by a period of refractoriness to short-term IFN stimulation (Borden and Murphy, 1971). The contraction of T^IFN^ in CFA and LPS treated mice (Fig. 5B and C), may reflect IFN refractoriness of naïve CD4^+^ T cells. Altogether, these data along with a recent study that showed IL-4 production during helminth infection transcriptionally alters naïve CD4^+^ T cells, demonstrate that even after removal, cytokine may have long-term impacts on the naïve CD4^+^ T cell state or responsiveness to stimuli (Even et al., 2024). These observations highlight the importance of considering naïve CD4^+^ T cell states during inflammatory diseases such as autoimmunity, infection and cancer, as aberrant naïve CD4^+^ T cells may contribute to increased disease burden, or exhibit varying responses to therapies due to the experience of inflammation.

### Human naïve CD4^+^ T cells display similar heterogeneity as mouse naïve CD4^+^ T cells

To determine if similar clusters were observed in humans, we performed scRNAseq on naïve CD4^+^ T cells isolated from PBMC of 6 healthy adult female donors, denoted as the DHMC cohort. CD4^+^ T cells were bead-enriched directly from whole blood and then stringently electronically sorted for naïve CD4^+^ T cells (singlet, viability dye-, CD4+, CD8-, CD45RA+, CD45RO-, CD127hi, CD25-, CD95-). We added to this population by computationally extracting naïve CD4^+^ T cells from two publicly available human healthy donor PBMC datasets. The first dataset, denoted Blood, included naïve CD4^+^ T cells filtered from the Tabula Sapiens dataset (Tabula Sapiens Consortium* et al., 2022). The second dataset included filtered naïve CD4^+^ T cells from only healthy controls in a previously published lupus dataset, with groups denoted as aHD for adult and cHD for pediatric healthy controls (Nehar-Belaid et al., 2020). Naïve CD4^+^ T cells were identified in these datasets by using the SingleR package to reference against the Celldex reference dataset DatabaseImmuneCellExpressionData (DICED) (Fig S15-17). Cells that were referenced as naïve CD4^+^ T cells were filtered and re-clustered. FACS sorted and public referenced cells were integrated with Seurat, SCT transformed, and co-clustered. Computationally extracted cells expressed canonical naïve marker genes at similar levels and occupied the same dimensional space after PCA reduction and UMAP projection, compared with in-house sorted cells (Fig. 6A).

**Figure 6.**
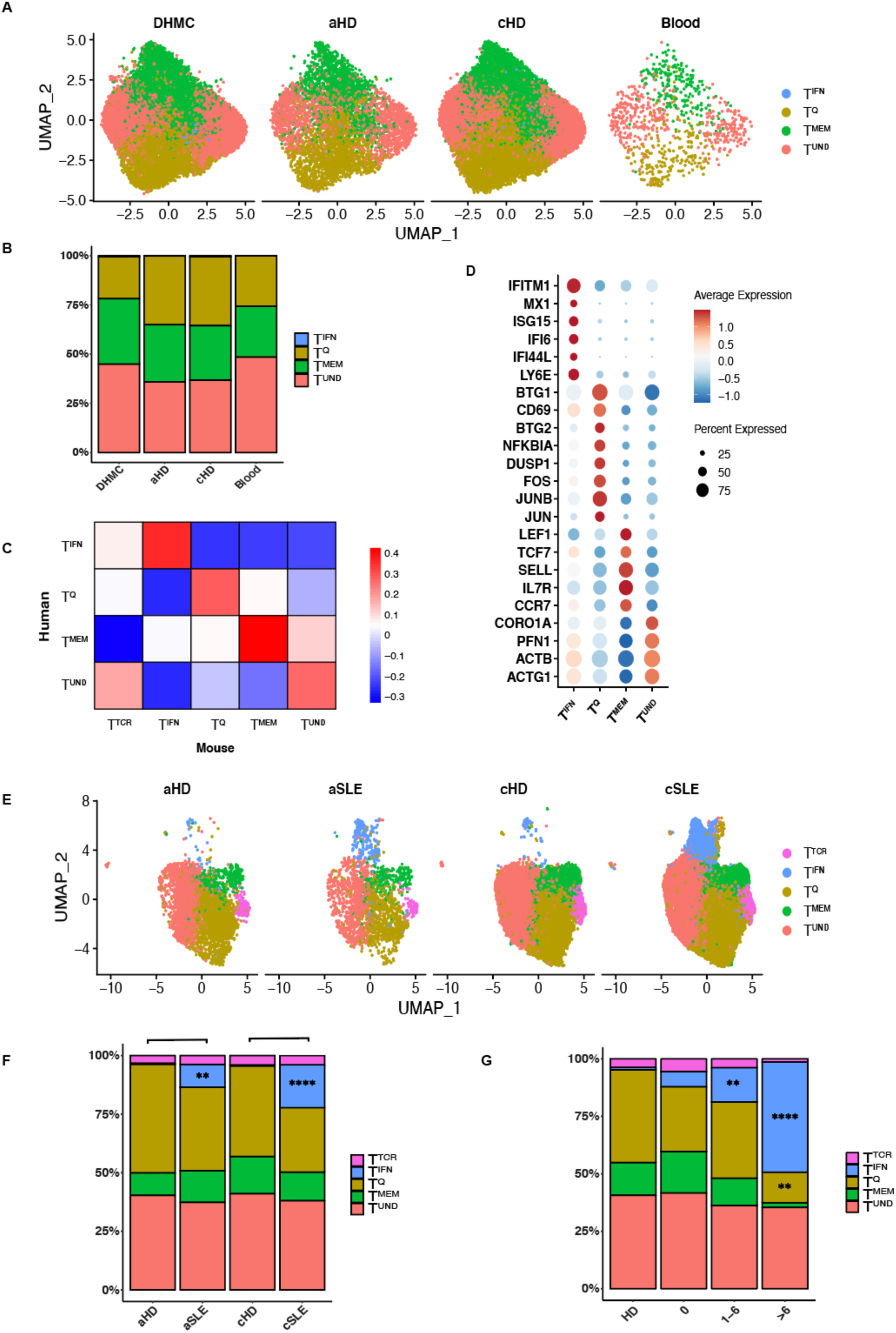
Human naïve CD4^+^ T cells during health and Systemic Lupus Erythematosus (SLE), in pediatric and adult donors. **A:** UMAP of naïve CD4^+^ T cells isolated from human PBMCs. In-house collected, sorted and sequenced patient cohort – DHMC (n=6), adult healthy controls from lupus cohort – aHD (n=5), pediatric healthy controls from lupus cohort – cHD (n=11) and Tabula Sapiens cohort – Blood (n=3). **B:** Stacked bar plot of frequencies in (A). **C:** Correlation plot of human and mouse naïve CD4^+^ T cell subset markers. Correlation values range between −1 and 1, with numbers above zero showing positive correlation. **D:** Dot plot showing marker genes of naïve CD4^+^ T cells from human blood in (A). **E:** UMAP of naïve CD4^+^ T cells from lupus cohort of human adult (aSLE) and pediatric (cSLE) lupus and healthy controls (aHD and cHD). **F:** Stacked bar plot of cluster frequencies in (E). All statistical significance testing was performed using 2-way ANOVA. * <0.05, ** <0.01, ***<0.001. **G:** Stacked bar plot of cluster frequencies by disease score. (Patient and donor data in this figure includes aHD: n=5, aSLE: n=4, cHD: n=11, cSLE: n=27). All statistical significance testing was performed using 2-way ANOVA. * <0.05, ** <0.01, ***<0.001.

In each of these subsets of healthy human naïve CD4s, we also identified 4 clusters: T^IFN^, T^Q^, T^MEM^ and T^UND^ that shared transcriptional features with their mouse cluster counterparts (Fig. 6A). In support of the reliability of the extracted naïve CD4^+^ T cells, we found all four clusters in human naïve CD4^+^ T cells that were observed in murine naïve CD4^+^ T cells, across 3 healthy human datasets (Fig. 6B). In line with murine blood naïve subsets, T^TCR^ was also not observed in naïve CD4^+^ T cell clusters in human blood (Fig. 6A and 4B). This may be due to lowered interactions with MHCs on APCs in the blood. In order to compare the clusters between the healthy human and C57BL/6J mouse naïve datasets, the Pearson correlation score was calculated between the normalized variable genes of each dataset, with zero representing no correlation and values above zero showing positive correlation. Human clusters showed moderate correlation (scores between 0.3 to 0.4) with the corresponding murine counterparts (Fig 6C). Dot plots show comparable marker genes in human and murine naïve CD4^+^ T cell clusters (Fig 6D). Genes shared between mouse and human T^IFN^ included *Isg15, Bst2, Irf7, Stat1, Isg20* and *Ly6e*. Genes present in murine T^IFN^ but not in human T^IFN^ included *Ifit3, Ifit1, Ifi47* and *Oasl2*, but *IRF9* was enriched in human but not mouse. For T^UND^, shared cluster markers were *Actb* and *Actg1*. In T^Q^, the shared markers were *Jun, Junb, Fos* and *Dusp1* with mouse T^Q^ expressing more *Jund* and *Klf6* than human T^Q^, and human T^Q^ expressing *Cxcr4*. T^MEM^ shared genes were *Il7r* and *Sell*. Mouse T^MEM^ had higher *Tcf7, Klf3* and *Itga4* expression. These data support the human relevance of naïve CD4^+^ T cell heterogeneity, and highlight the evolutionary importance of conserving naïve T cell transcriptional states.

### Human naïve CD4^+^ T cells in autoimmune disease

Lupus has been established as a multifactorial disease involving heightened autoantibody and IFNα profiles (Caielli et al., 2023). However, prior studies of lupus have not documented the impact of disease on the naïve T cell transcriptional profile, or its potential impact on the functional bias imposed. This is of importance in the clinic, as altered resting T cell heterogeneity and function could influence disease outcomes. To evaluate the potential impact of autoimmune disease on human naïve CD4^+^ T cell transcriptional heterogeneity, we analyzed filtered naïve CD4^+^ T cells from 4 adult (aSLE) and 27 pediatric (pSLE) lupus patients and matched healthy controls (5 aHD and 11 cHD) (Fig. 6E) (Nehar-Belaid et al., 2020). In both adult and pediatric lupus, the frequency of T^IFN^ was increased relative to healthy controls; from 0.56% in aHD to 9.74% in aSLE, and 0.54 in cHD to 18.35% in cSLE (Fig. 6E and 6F). No clusters were significantly different between adult healthy and pediatric healthy controls. The T^IFN^ cluster was statistically increased in pediatric lupus patients compared to healthy pediatric donors (p <0.0001). Furthermore, the T^IFN^ cluster size increased as a function of worsening lupus disease score, measured using SLEDAI (Fig. 6G). In healthy controls, the T^IFN^ cluster is at a frequency of 1.0%. For lupus patients with SLEDAI disease score of 0, the T^IFN^ is 6.7%, for a disease score between 1-6 T^IFN^ is 15% and for a disease score higher than 6 T^IFN^ climbs up to 48% (Fig. 6G). The r value 0.52 for the correlation between T^IFN^ and SLEDAI disease score and p value is 0.0002. This demonstrates that naïve CD4^+^ T cells are impacted and suggests that this increase may be mediated by the increase in environmental IFNα during lupus disease.

Therefore, the increase in T^IFN^ during lupus, and with worsening disease burden as measured by the SLEDAI score, demonstrates that the naïve CD4^+^ T cell compartment is responsive to systemic inflammation. Whether the altered naïve T cell heterogeneity influences the trajectory of the disease is yet to be explored.

## Discussion

In this study, we establish transcriptional and functional heterogeneity of the resting CD4^+^ T cell compartment. Our single cell transcriptional profiling revealed 5 subsets in the naïve CD4^+^ T cell compartment: a quiescence cluster (T^Q^), a memory-like cluster (T^MEM^), a TCR reactive cluster (T^TCR^), an IFN responsive cluster (T^IFN^) and an undifferentiated cluster (T^UND^). We observed similar naïve CD4^+^ T cell clusters in humans and in mice (Fig. 1 and Fig. 6). The observed naïve CD4^+^ T cell transcriptional heterogeneity impacts on the later CD4^+^ T cell differentiation trajectories.

We have identified functional biases in each of the resting T cell subsets. We enriched for naïve CD4^+^ T cell subsets for functional analyses using TF reporter mice. These enriched naïve T cell subpopulations may not comprehensively represent the subsets defined by scRNAseq, but clearly establish the existence of functional heterogeneity amongst resting T cells. We demonstrate that the Nur77^HI^ naïve CD4^+^ T cell subset is hyporesponsive to activation, and produces less cytokine upon activation, as suggested by prior studies (Zinzow-Kramer et al., 2019; Sood et al., 2019). Our data demonstrates naïve T cells with a heightened Mx1-GFP profile skew towards Th1 and Th17 lineages, and away from a Th2 trajectory (Fig. 1G). The heightened expression of ISGs may enhance the readiness or “preparedness” of naïve T cells, through the induction of high numbers of idling ribosomes in naïve T cells (Wolf et al., 2020). T^Q^ also skewed towards Th1 and Th17 lineages and away from a Th2 trajectory (Fig. 1G). While *Btg1* and *Btg2* are classic quiescence factors, KLF2 has an additional has a role in mediating thymic egress and T cell migration (Hwang et al., 2020; Carlson et al., 2006). KLF2 has been shown to modulate S1PR, CD62L and β7 integrin expression, all vital to T cell circulation and homing (Carlson et al., 2006). Transcriptional changes in KLF2 expression, and consequently the proportion of the T^Q^ cluster between the spleen and blood, may be an additional dynamic force that allows constant circulation of naïve T cells between lymphoid organs.

The ontogeny of these naïve CD4^+^ T cell clusters was also characterized in this study. Naïve CD4^+^ T cell transcriptional heterogeneity is established in the thymus, as T cells enter the SP stage (Fig 2). It appears that TCR specificity plays a minor role in cluster designation as clusters in TCR Tg mice are very similar to those found in WT mice (Fig. S11). Because these naïve CD4^+^ T cell clusters are largely maintained in the periphery, even under immune duress, the clusters imposed in the thymus are quite rigid. It is striking that under conditions where the immune context has been changed demonstrably – such as in germ free or CFA/LPS treated mice – the composition of the naïve T cell clusters is unchanged. This demonstrates the rigidity and prolonged maintenance of the cluster frequencies induced in the thymus. Exactly what cells or mediators in the thymus control cluster designation is yet to be resolved.

We have identified some immune cues that can impact on cluster heterogeneity. We report that the T^TCR^ cluster of naïve CD4^+^ T cells is lost upon blocking of tonic signaling (Fig. 3). In addition, blocking tonic signaling results in the downregulation of TCR signaling genes across all clusters. Notably, while cytokines are not classically accepted as modulators of the naïve CD4^+^ T cell state, IFNβ treatment expanded T^IFN^, demonstrating that cytokines can influence naïve T cell transcriptional states (Fig. 5E). An in vivo study where mice were treated with 86 cytokines reported that immune cells experience cytokine-driven cellular polarization, as determined by scRNAseq (Cui et al., 2024). In a published examination of human naïve CD4^+^ T cells pre-treated with IL-12 before in vitro activation, the authors observed increased phosphorylation of TCR signaling tyrosine kinase Lck, TCR induced cytokine production and oxidative metabolism following TCR stimulation (Vacaflores et al., 2016). Furthermore, we found that inflammation also alters the naïve CD4^+^ T cell state. Naïve CD4^+^ T cells from CFA treated mice were less responsive to activation and differentiation in multiple T helper cell skewing conditions than naïve CD4^+^ T cells from untreated mice (Fig. 5D). Hence, exposure of resting T cells to immune stimuli results in a subsequent altered activation response, even if they remain in their naïve T cell state at first. This phenomenon could be compared to “trained immunity,” defined in innate cells as when environmental exposures establish innate immune memory (Netea et al., 2020).

To further explore how the naïve CD4^+^ T cell state changes in humans in the presence of inflammation, we assessed naïve CD4^+^ T cell clusters in human autoimmune disease. During lupus progression, naïve CD4^+^ T cells from both adult and pediatric patients exhibited an expanded T^IFN^ cluster, which increases as lupus disease score increases. Lupus, a systemic autoimmune disease that is primarily characterized by autoantibodies, has been shown to have an increased IFN signature in multiple immune cells (Postal et al., 2020). The increased IFN signature in naïve CD4^+^ T cells demonstrates that naïve T cells are not just bystander cells; they respond to their environment through altered genetic expression. Moreover, altered epigenetic programing in naïve CD4^+^ T cells from lupus patients has been reported, whereby hypomethylation of interferon-regulated genes results in increased transcriptional accessibility of IFN genes that promote disease (Coit et al., 2013). Of note, IFN refractoriness has been described as a period of exhaustion and hyporesponsiveness to IFN signaling (Borden and Murphy, 1971). Whether this skewed naïve T cell state contributes to disease pathogenesis because of altered functional status is yet to be determined.

In conclusion, the resting CD4^+^ T cell compartment is heterogeneous in transcriptional profile and in functional bias. Each of the CD4^+^ T cell clusters appear subject to environmental influences that change their transcriptional profiles, and in some cases their functional trajectory upon activation. The finding that immune context alters the subsequent functional immune responses of resting T cells is a model for “trained T cell immunity”. Most critical is determining how trained T cell immunity in human autoimmune diseases alters the progression of those diseases.

## Materials and Methods

### Mice

C57BL/6J (JAX strain 000664; B6), B6.Cg-Tg(TcraTcrb)425Cbn/J (JAX strain 004194; OT-II), B6.Cg-Ptprca Pepcb Tg(TcrLCMV)1Aox/PpmJ (JAX strain 030450; SMARTA), B6.SJL-*Ptprc^a^ Pepc^b^*/BoyJ (JAX strain 002014; CD45.1), C57BL/6-Tg(Nr4a1-EGFP/cre)820Khog/J (JAX strain 016617; Nur-77-GFP), B6.Cg-*Mx1^tm1.1Agsa^*/J (JAX strain 033219; Mx1-GFP), mice were purchased from the Jackson Laboratory (Bar Harbor, Maine). Germ-free and matched specific-pathogen free C57BL/6NCrl (NCI strain 556; B6) mice were purchased from Charles River (Wilmington, MA), and were euthanized for analysis on arrival. OT-II x *Rag2^em3Lutzy^*/J (OT-II crossed with JAX strain 033526; OT-II x Rag2-/-), C57BL/6-Tg(Rag2-EGFP)1Mnz (JAX strain 005688 backcrossed to B6; Rag2-GFP), and B6(C)-*Klf2^tm1.1Khog^*/JmsnJ (JAX stock #036331; KLF2-GFP) mice were generously provided by Kris Hogquist and Stephen Jameson at the University of Minnesota, and were described previously (Weinreich et al., 2009). Deletion of VISTA in CD4^+^ T cells was performed by crossing VISTA^fl/fl^ mice to hemizygous B6.Cg-Tg(Cd4-cre)1Cwi/Bfluz (JAX stock #022071), which were generously provided by Sam Lee at Yale University (Yoon et al., 2015). Cre-littermate controls were used for thymic DP, thymic CD4^+^ SP and splenic naïve CD4^+^ T cell groups in Fig. 2A. Mouse strains were maintained in a pathogen-free facility at the Geisel School of Medicine at Dartmouth College, NH, USA. All mouse studies described in this work were carried out in accordance with the principles of the Guide for the Care and Use of Animals and were approved by the Institutional Animal Care and Use Committee of Dartmouth College, NH, USA (Protocol 2012).

### Naïve CD4^+^ T cell isolation

#### Murine tissue processing

Spleens, lymph nodes or thymii were dissociated in HBSS between the frosted sections of glass slides, and were then filtered through 70nm cell strainers (Becton Dickinson, Franklin Lakes, NJ).

#### Murine blood processing

Blood was collected by cardiac puncture into 20ul of heparin (Selleckchem; Houston, TX). Red blood cells were lysed with Biolegend RBC lysis buffer according to manufacturer’s directions.

#### Human and Murine Naïve CD4^+^ T cell pre-enrichment

Mouse cells were first pre-enriched by negative selection, using the EasySep™ Mouse Naïve CD4^+^ T Cell Isolation Kit according to manufacturer instructions (StemCell, Vancouver, BC). Isolated cells were stained and FACS sorted for naïve CD4^+^ T cells (singlet, viability dye-, CD4^+^, CD8^-^, CD44^lo^, CD62L^+^, CD25^-^, CXCR3^-^).

PBMCs were collected from healthy adult female donors. Human PBMC were pre-enriched directly from total blood with Biolegend EasySep™ Direct Human CD4^+^ T Cell Isolation Kits according to manufacturer’s instructions, and then stringently electronically sorted for naïve CD4^+^ T cells (singlet, viability dye-, CD4^+^, CD8^-^, CD45RA^+^, CD45RO^-^, CD127^hi^, CD25^-^, CD95^-^).

#### Antibodies and sorting

Antibodies used for sorting murine cells were CD4 BV510 (RM4-5), CD8 PerCP/Cy5.5 (53-6.7), CD44 PE (IM7), CD62L bv421 (MEL-14), CD25 PE/Cy7 (PC61) and CXCR3 APC/Cy7 (CXCR3-173). Antibodies used for sorting human cells were CD127 BV421, BV510 CD95, CD45RA APC, CD8 APC-CY7, CD4 AF488, PerCP/Cyanine5.5 anti-human CD45RO and CD25 PECy7. Dead cells were labeled with Live/Dead Fixable Near-IR dead cell stain (Invitrogen; catalog no. L10119) in PBS. Cells were washed and then flow-sorted using FACS-ARIA II (BD Biosciences). Purity was validated by using subsequent flow cytometry. For enrichment of GFP reporter expression, the top 10-15% and bottom 10-15% of GFP expressing naïve CD4^+^ T cells were sorted from Nur77-GFP and KLF2-GFP mice. For Mx1-GFP mice, positive GFP expression was identified using a gate set by Wild-Type mice (which approximated to the top 1-2% of GFP expression in the Mx1-GFP naïve CD4^+^ T cells) while GFP negative was isolated using a gate set on the bottom 10% of GFP expression in the Mx1-GFP naïve CD4^+^ T cells.

### Single cell RNA sequencing and processing

#### Cell hashtagging for scRNAseq experiments

TotalSeq hashtag antibodies were used to allow multiplexing of up to 6 samples per lane during scRNAseq processing. Each sample to be pooled was stained with 2ug of a differently barcoded Biolegend TotalSeq hashtag antibody (Hashtags 1 to 6 were used) along with the cocktail of fluorescently labeled antibodies as described above. After staining, cells were then washed 3 times with 3.5ml of PBS with 1% BSA and then mixed together prior to sorting.

#### CITE-Seq sample preparation

Up to 2 million hashtagged and fluorescently sorted naïve CD4^+^ T cells were stained with Mouse or Human TotalSeq™-C Universal Cocktail V1.0 CITEseq antibodies (Biolegend) according to manufacturer directions. Cells were then washed 3 times with 3.5ml of PBS with 1% BSA prior to processing for scRNAseq.

#### Droplet-based 5ʹ end single-cell RNA sequencing

Droplet-based 5ʹ end (scRNAseq) was performed by the 10x Genomics platform and libraries were prepared by the Chromium Single Cell NextGEM 5’ v2 chemistry Reagent kit (CG000331, 10x Genomics, CA, USA) and sequenced on a NextSeq2000 using 28×90bp reads for GEX libraries according to manufacturer’s protocol (10x Genomics, CA, USA). 16,500 cells were loaded onto the 10x chip to target 10,000 sequenced cells, according to manufacturer instructions. The CellRanger Single-Cell Software Suite (10x Genomics) versions v2.1.1 to v7.1.0 were used to perform barcode processing and transcript counting after alignment to the mm10 reference genome with default parameters. Sample barcode hashtags were matched to cells using the HTODemux function of the Seurat package. Droplets containing reads from multiple hashtags were deemed doublets or triplets and filtered out. Hashtags associated with low numbers of reads, likely suggestive of free-floating RNA, were also removed from consideration. Quality control was performed using the Seurat pipeline, with only cells that had between 200 and 2200 genes used for further analyses. Cells with more than 5% mitochondrial gene content were also filtered out to omit dying cells with aberrant RNA presence. TCRa and TCRb genes were removed from consideration as variable features. Clusters with cells expressing very high levels of mitochondrial genes, *Foxp3,* or *Cxcr3* were removed from the dataset to filter out Tregs and stem cell memory T cells. Normalization was performed using the SCTransform function of the Seurat package.

#### Integration and batch correction between datasets

In order to standardize and allow identification of the T^Q^ cluster, all datasets generated in this study were integrated with our previously published dataset for naïve CD4^+^ T cells from VISTA KO and WT mice (ElTanbouly et al., 2020), available under BioProject accession number PRJNA587711, as well as naïve CD4^+^ T cells from multiple tissues (spleen, lymph node and blood) from C57BL/6J mice.

Data from scRNAseq replicates were integrated using Seurat integration with Canonical Correlation Analysis (CCA) using the Seurat R package (v.4.1.2). Code used followed the workflow described in the Satija lab vignette “Performing integration on datasets normalized with SCTransform” at https://github.com/satijalab/seurat/blob/HEAD/vignettes/integration_introduction.Rmd

To assess whether batch effects were present in the data after integration UMAPs before and after integration were visually inspected.

#### Dimension reduction and clustering

The RunPCA function was used to perform principal component analyses followed by the RunUMAP and FindNeighbors functions of the Seurat package. The top 3000 genes were selected for integration, and the top 30 principal components were used to generate the UMAPs. The FindClusters function was used to cluster by Louvain clustering. The FindAllMarkers function (Seurat v.4.1.2) was used to identify marker genes for each cluster using the Wilcoxon Rank Sum, MAST and Roc tests. Marker genes were visually evaluated by DotPlot function (Seurat v.4.1.2) and were validated using the Single-Cell Clustering Assessment Framework for AUC scores of > 0.7.

#### Pathway analysis and scoring of biological processes

To compare between groups, ranked lists of Log2 gene expression comparison between groups were utilized for pathway expression analysis using the Broad Institute Gene Set enrichment analysis (GSEA; v4.3.2). The C7 (immunesigdb.v2023.1) and C2 (v2023.1) pathway gene sets were tested for potential enrichment of pathway gene in each cluster. Pathways with Benjamini–Hochberg–adjusted *P* values < 0.01 were considered significant.

#### RNA velocity analysis

BAM files from the cell ranger output for naïve CD4^+^ T cell scRNAseq from the spleen were used to create loom files. The scVelo toolkit was used in Python to generate RNA velocity calculations, with code adapted from https://smorabit.github.io/tutorials/8_velocyto/, which were then projected onto UMAPs created by the Seurat pipeline in R.

#### Public scRNAseq data

Naïve CD4^+^ T cells were extracted from Tabula Sapiens (Tabula Sapiens Consortium* et al., 2022) and lupus public scRNAseq datasets (Nehar-Belaid et al., 2020) using the ‘SingleR’ (v.2.2) package to annotate cell identities against the built-in Celldex reference dataset Database of Immune Cell Expression (DICE) dataset (Aran et al., 2019; Schmiedel et al., 2018). DICE contains the normalized expression values of 1561 bulk RNA-seq samples of sorted cell populations. Cells referenced as naïve CD4^+^ T cells were filtered out and reclustered to identify human naïve CD4^+^ subsets.

#### Calculation of correlation scores

In order to compare the clusters between the healthy human and autoimmune mouse naïve datasets, a correlation score was calculated. First, gene-wise normalization was performed by calculating the z-score of each gene on the log transformed gene count. Genes were filtered for the genes that defined cell types and subsets. Next, the Pearson correlation coefficient was calculated using the cor() function in R.

### Bulk RNA sequencing and processing

Naïve CD4^+^ T cells (isolated as described above) from Mx1-GFP mice were sorted on the Mx1-GFP+, with a gate set using WT mice, which was the top 1% of cells. The Mx1-GFP-population was sorted using a gate set on the bottom 10-15% of naïve CD4^+^ T cells. Nur77-GFP and KLF2-GFP were used to isolate the self-reactive and quiescent populations respectively. Since all naïve CD4s express Nur77 and KLF2, just to different degrees, the top and bottom 10-15% of naïve CD4^+^ T cells were sorted for these strains. Samples were sorted into buffer RTL. Then, RNA was isolated and prepped for sequencing using the Takara Pico v3 kit.

### Mouse treatments

For testing the impact of interferon treatment, mice were injected with 0.25ug of human IFNb (Peprotech) i.p. daily for 5 days, or daily for 5 days followed by 5 days of rest. Treated mice were compared to untreated controls. All mice were analyzed together 30 minutes after the last IFNb injection.

For testing the impact of tonic TCR signaling, mice were injected i.p. for 3 consecutive days with 1mg of either anti-MHCII (anti-I–Ab, clone Y-3P; BioXCell, Lebanon NH) or isotype control (clone C1.18.4; BioXCell). Mice were analyzed one day after the final antibody injection.

For testing the impact of an inflammatory environment, mice were injected i.p. with a 1:1 emulsion of 100ug CFA and PBS or 10ug LPS in PBS. Mice were analyzed or sorted as above for functional skewing experiments 7 days after treatment.

For the tissue transfer experiment, blood was collected from CD45.1 mice, and red blood cells depleted using RBC lysis buffer and followed by a spin down. The remaining lymphocytes were transferred into CD45.2 mice via an i.v. injection. 3 days later, spleens and blood were collected from the mice, stained with the naïve panel and sorted as above. Both CD45.1 and CD45.2 naïve CD4^+^ T cells were collected. Single cell RNA sequencing was performed on the sorted naïve CD4^+^ T cells.

### Functional studies

GFP reporters were used to isolate subsets of naïve CD4^+^ T cells for in vitro functional experiments as above. 25,000 naïve CD4^+^ T cells were cultured with 100,00 irradiated splenocytes (irradiated at a dose of 50 Gray) in T helper cell skewing conditions for 5 days. Plates were coated a day prior with anti-CD3 and anti-CD28 at 1 ug/ml for all culture conditions except for Th17, which had 2 ug/ml anti-CD3 and 1 ug/ml anti-CD8. The culture conditions included 30 U/ml IL-2 for the Th0 control, 30 U/ml IL-2, 15 ng/ml IL-12 and 5 ug/ml aIL-4 clone 11B11 for the Th1 skewing, 30 U/ml IL-2, 10 ng/ml IL-4 and 5 ug/ml aIFNγ (XMG1.2) for Th2 skewing, and 20 ng/ml IL-6, 3 ng/ml MsTGFβ, 10 ng/ml IL-1b, 20 ng/ml IL-23, 5 ug/ml anti-IFNγ (XMG1.2) and 5 ug/ml anti-IL-4 (11B11) for Th17 skewing (Flaherty and Reynolds, 2015). IL-12p70 PeproTech cat# 210-12; IL-6 PeproTech cat# 216-16; IL-4 PeproTech cat# 214-14; TGFβ1 R&D cat# 7666-MB-005; aIL-4 (clone 11B11) BioLegend cat# 504122; aIFNγ (clone XMG1.2) BioLegend 505834; IL-1b PeproTech cat# AF-211-11b; IL-23 R&D cat# 1887-ML-010. Supernatant was collected after 5 days and Luminex was performed to identify cytokine production.

### Statistics

For the functional studies, z scores were calculated to plot Luminex data on heat maps. Multiple unpaired t-tests were used to determine significance. Percent change comparing reporter GFP+ increase from GFP-, or CFA treated increase from untreated controls was calculated to demonstrate magnitude of change in cytokine production. To determine if cluster frequency changes were significant, multiple unpaired t-tests or two-way ANOVA tests were performed for each cluster grouping as appropriate.

### Naïve CD4 cluster names

Quiescence cluster: T^Q^

Memory-like genes in naïve CD4s cluster: T^MEM^

TCR signaling/self-reactive/tonic signaling cluster: T^TCR^

Interferon-Stimulated Genes (ISGs) cluster: T^IFN^

Undifferentiated cluster: T^UND^

## Acknowledgments

We thank Kris Hogquist and Stephen Jameson at the University of Minnesota for their insightful comments during the conception and development of this manuscript. Flow cytometry and flow sorting experiments were carried out by DartLab, the Immune Monitoring and Flow Cytometry Shared Resource at the Dartmouth Cancer Center, with NCI Cancer Center Support Grant 5P30 CA023108-41. RNA-sequencing experiments were carried out in the Genomics and Molecular Biology Shared Resource (RRID:SCR_021293) at Dartmouth which is supported by NCI Cancer Center Support Grant 5P30CA023108 and NIH S10 (1S10OD030242) awards. Single cell studies were conducted through the Dartmouth Center for Quantitative Biology in collaboration with the GMBSR with support from NIGMS (P20GM130454) and NIH S10 (S10OD025235) awards. Experimental outlines, graphical summary and Supplemental figures S21 and S22 were generated using BioRender.

## Author contributions

Conceptualization: R.J.N, A.S., M.J.T. and J.L.L; Methodology: R.J.N., A.S., M.J.T. and J.L.L; Investigation: A.S., J.L.L. M.E. and W.C.; Writing – initial manuscript: A.S. and J.L.L; Writing – reviewing & editing: R.J.N., A.S., J.L.L and M.J.T.; Computational analysis: A.S. and J.L.L.; Patient sample collection: C.B.; Resources: R.J.N.; Supervision: R.J.N. and M.J.T.

