## Supplemental Figures for "Heterogeneity and plasticity of the naïve CD4^+^ T cell compartment"

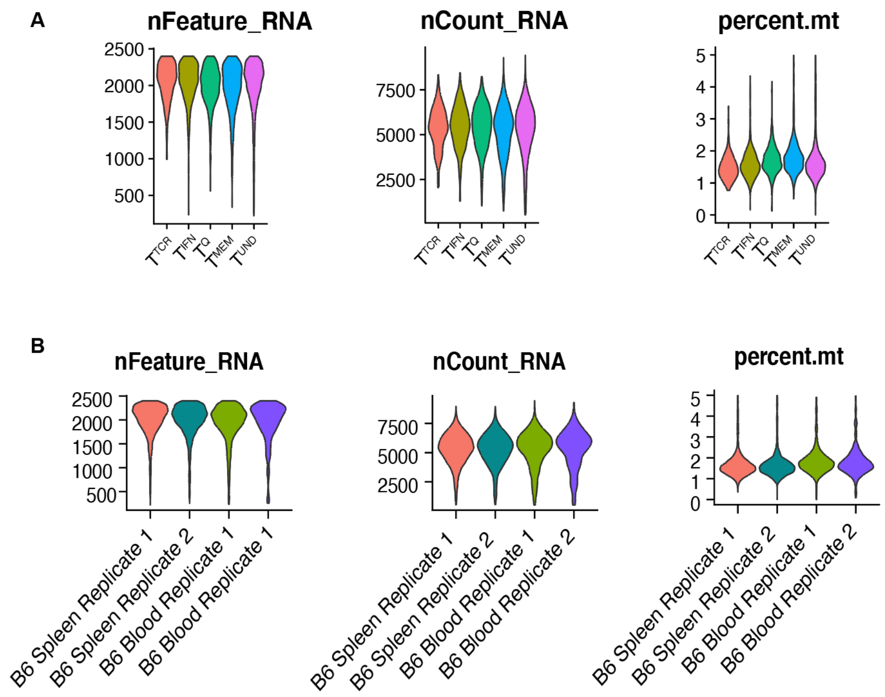


**Figure S1. Representative plots for mouse single cell quality control pipeline showing nFeature, nCount and mitochondrial content. A.** nFeature, nCount and mitochondrial content per cluster after quality control filtering. **B.** nFeature, nCount and mitochondrial content separated out by tissue and donor.

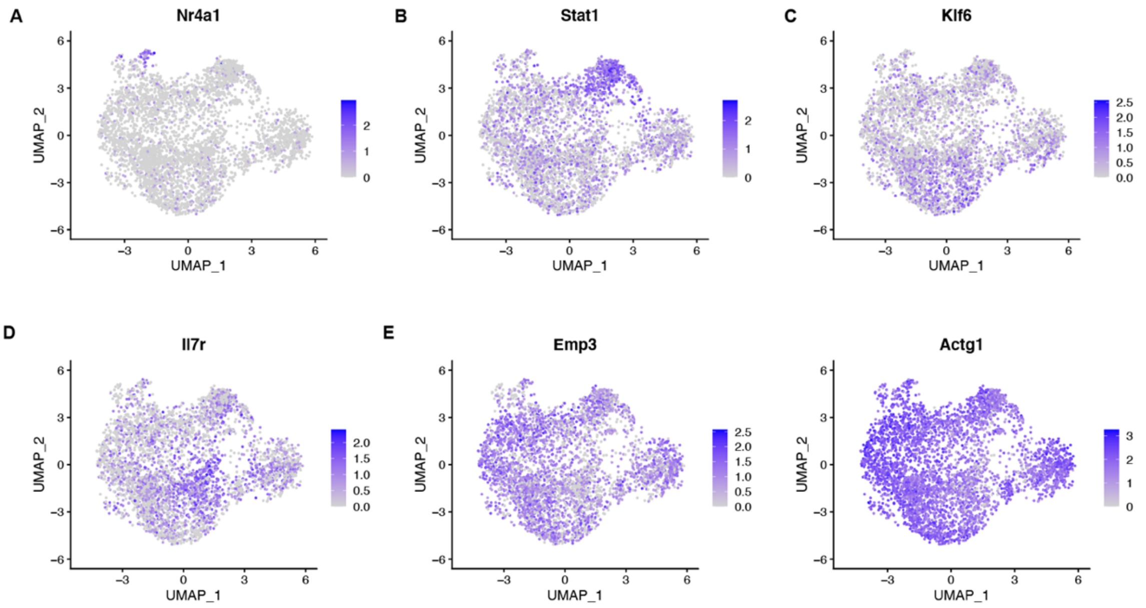


**Figure S2. Feature plots of marker genes plots for mouse single cell data.** Feature plot showing expression of *Nur77, Stat1, Klf6, Il7r, Emp3* and *Actg1* on UMAP.

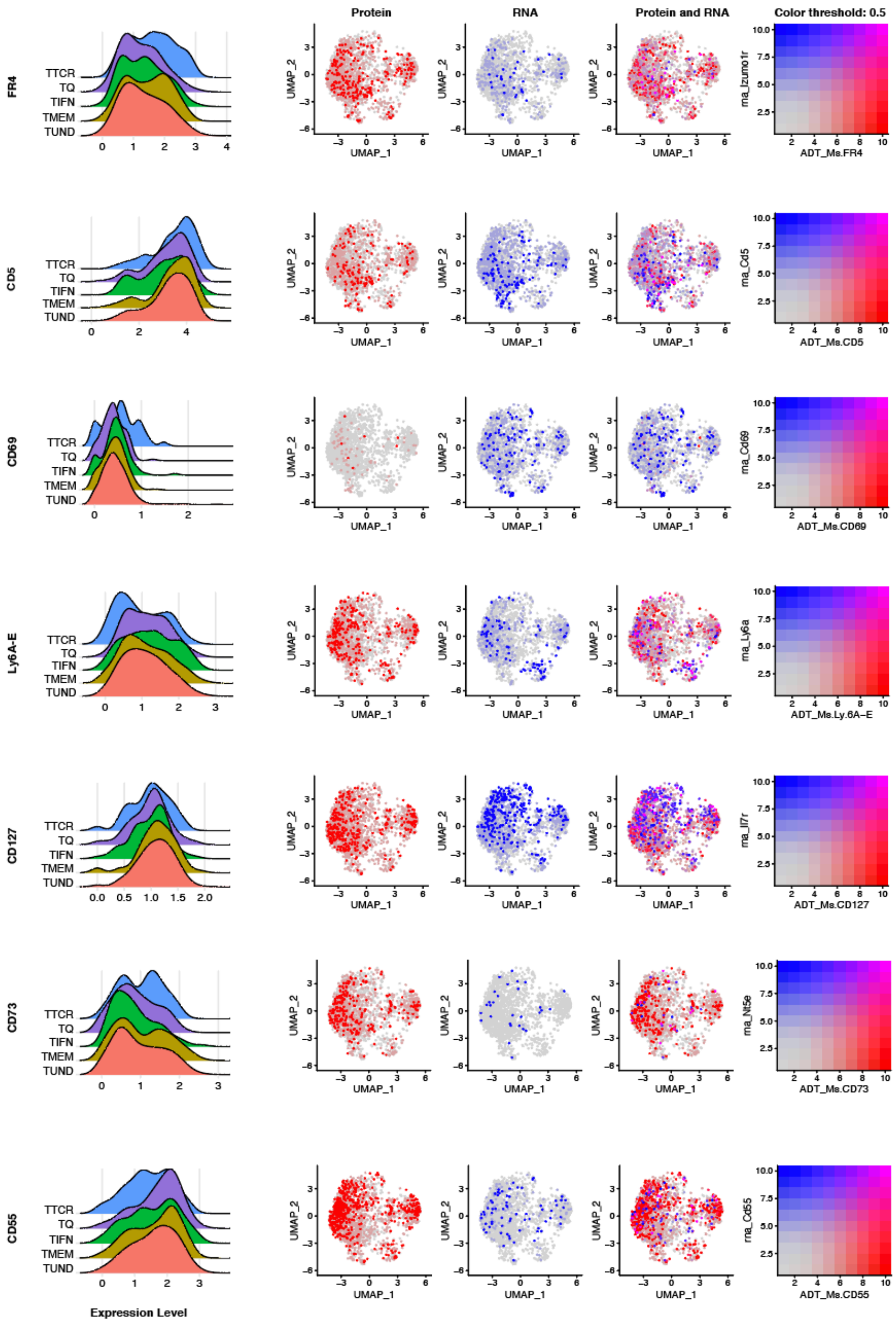

**Figure S3. CITE-seq of mouse naïve CD4$^{+}$ T cells does not show overlap of surface markers with transcriptional clusters.** CITE-Seq performed on murine naïve CD4$^{+}$ T cells isolated from C57BL/6J mice. FR4, CD5, CD69, Ly6A-E, CD127, CD73 and CD55 selected as representative markers of the CITE-Seq data. From left to right: Ridge plots showing expression of select protein markers by clusters ($T^{TCR}$ in blue, $T^{Q}$ in purple, $T^{IFN}$ in green, $T^{MEM}$ in brown and $T^{UND}$ in red). Feature plots of RNA UMAP showing protein expression in red, RNA in blue and overlap of protein and RNA in purple.

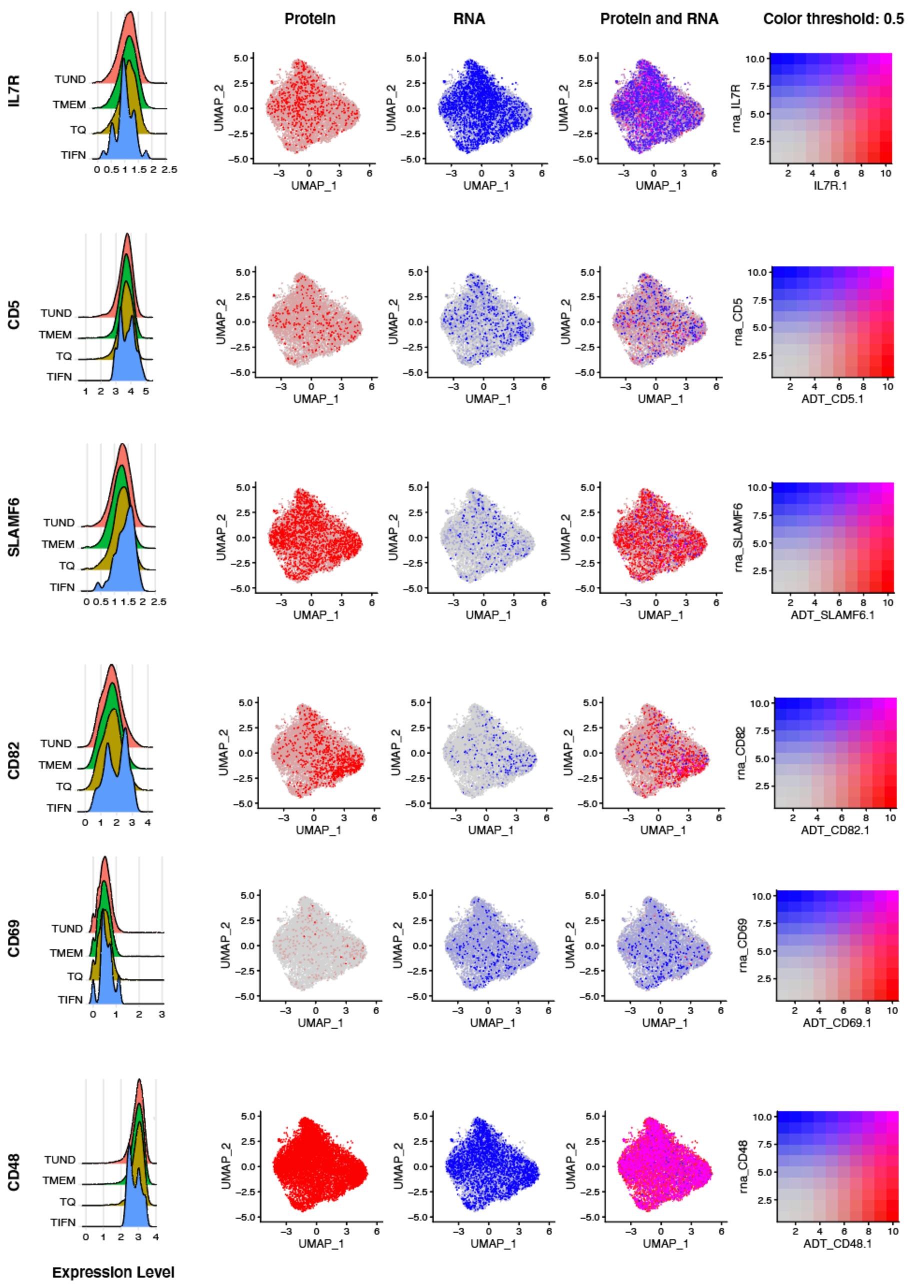


**Figure S4. CITE-seq of human naïve CD4⁺ T cells doesn't show overlap of surface markers with transcriptional clusters.** CITE-Seq performed on human naïve CD4⁺ T cells isolated from healthy donors. IL7R, CD5, SLAMF6, CD82, CD69 and CD48 selected as the best markers representative of the CITE-Seq data. From left to right: Ridge plots showing expression of select markers by cluster ($T^{Q}$ in brown, $T^{IFN}$ in blue, $T^{MEM}$ in green and $T^{UND}$ in red). Feature plots showing protein in red, RNA in blue and overlap of protein and RNA in purple.

**A** **KLF2-GFP**

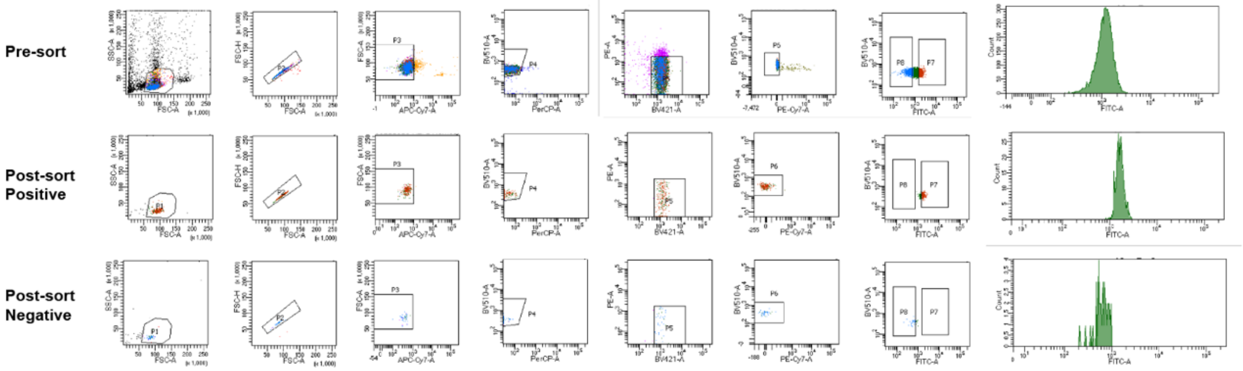


**B**

**Mx1-GFP**

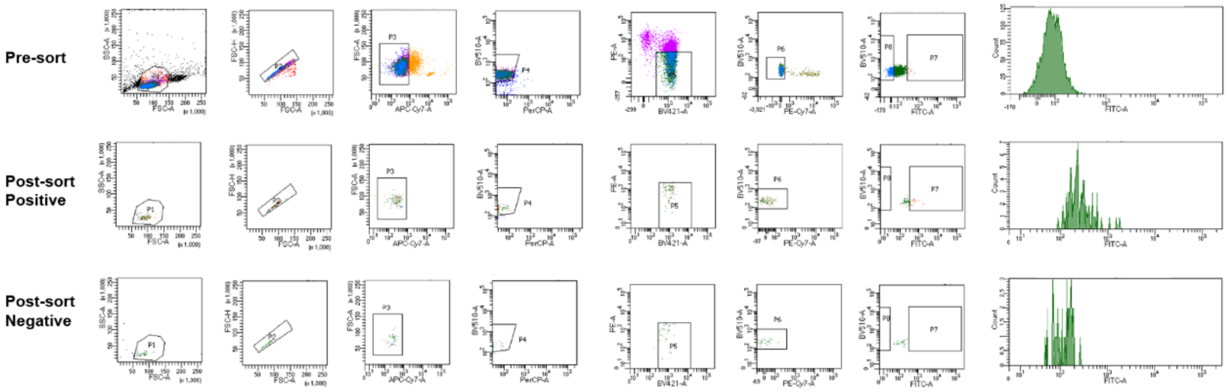

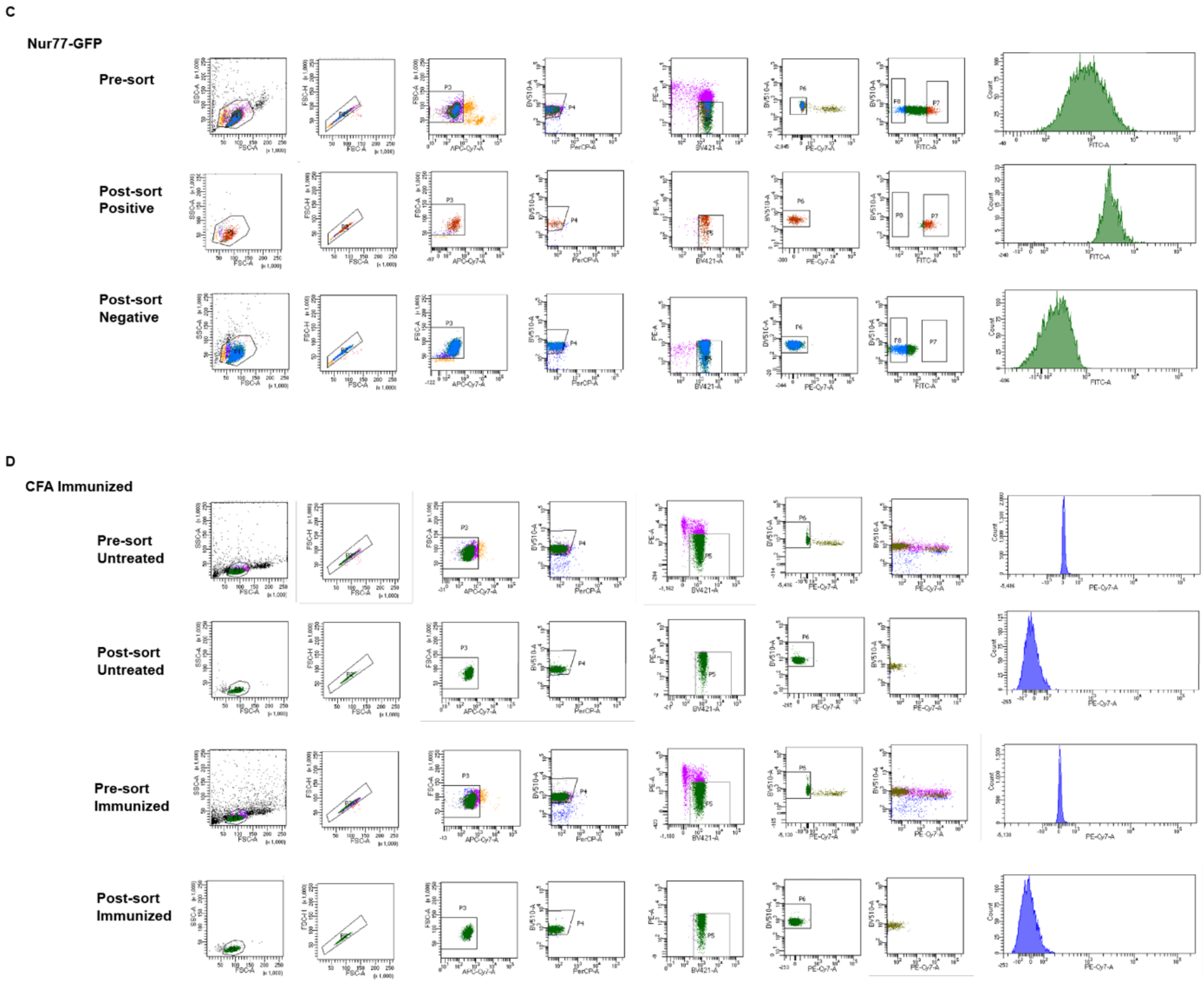


**Figure S5. Representative plots showing sort gating strategy, and pre and post enrichment flow plots for KLF2-GFP, Mx1-GFP, Nur77-GFP and CFA immunized GFP$^{HI}$, GFP$^{LO}$ and naïve CD4$^{+}$ T cells. A.** KLF2-GFP high vs low naïve CD4$^{+}$ T cell sort gating and purity check. **B.** Mx1-GFP high vs low naïve CD4$^{+}$ T cell sort gating and purity check. **C.** Nur77-GFP high vs low naïve CD4$^{+}$ T cell sort gating and purity check. **D.** CFA immunized naïve CD4$^{+}$ T cell sorting sort gating and purity check.

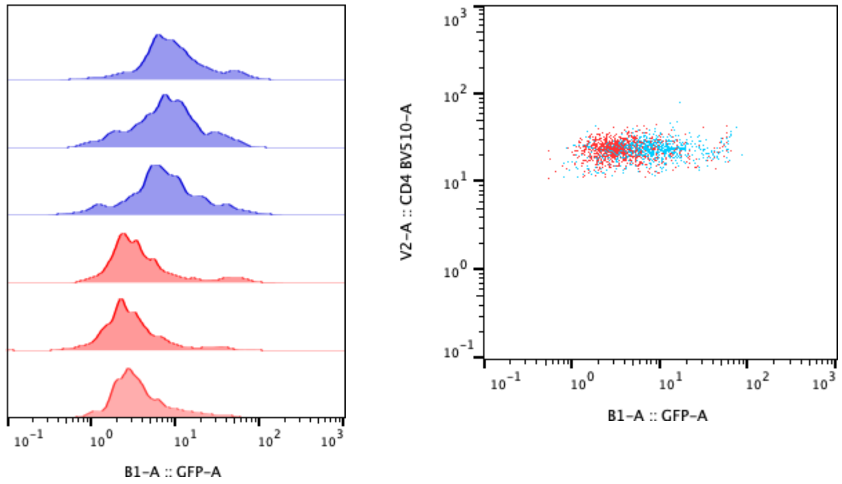


**Figure S6. Adoptive transfer of Nur77$^{HI}$ vs Nur77$^{LO}$ cells shows GFP reporting stability.** 300,000 naïve CD4s Nur77-GFP$^{HI}$ and Nur77-GFP$^{HI}$ (sorted using top 15% and bottom 15% GFP expression) were adoptive transferred into CD45.1 recipients. Nur77-GFP expression was assessed after 4 days. Blue = Nur77$^{HI}$ and red = Nur77$^{LO}$.

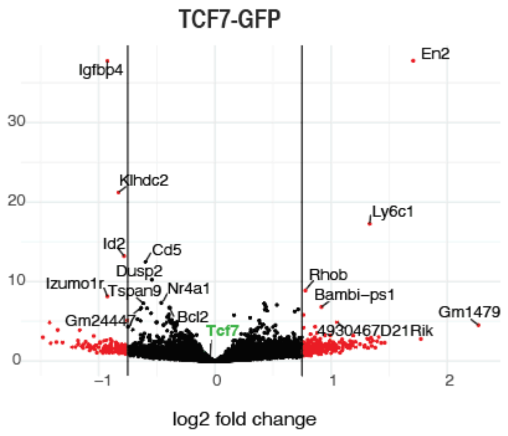


**Figure S7. Bulk RNA sequencing of naïve CD4$^{+}$ T cells from TCF7-GFP reporter mice shows no enrichment of T$^{MEM}$.** Volcano plots showing bulk sequencing of sorted of naïve CD4$^{+}$ T cells sorted from top and bottom 10-15% GFP reporter mice, corresponding to TCF7$^{HI}$ vs TCF7$^{LO}$ naïve CD4$^{+}$ T cells. The x axis is the log2fold change value and the y axis is -log10 of the p value. Genes labeled in green correspond to cluster markers. (n=4).

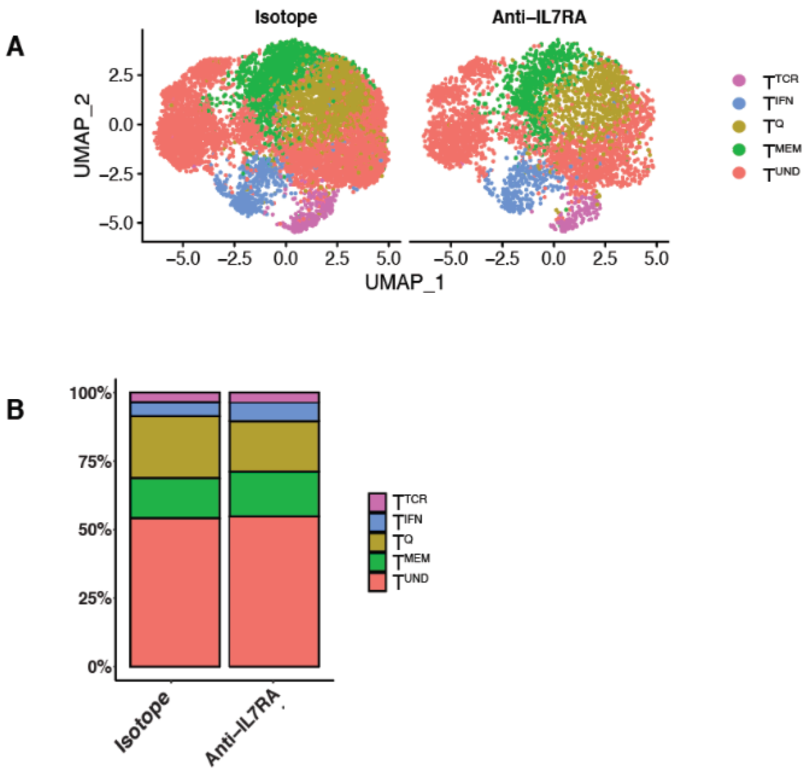


**Figure S8. Naïve CD4+ T cell clusters don't change in anti-IL7RA or isotype treated mice. A:** UMAP naïve $CD4^+$ in the spleens of anti-ILR7A or isotype treated mice. Mice were injected with 300ug/mouse anti-ILRA or isotype antibody every other day for a total of 3 treatments. **B:** Stacked bar plot of frequencies in (A), grouped by treatment. (n=2).

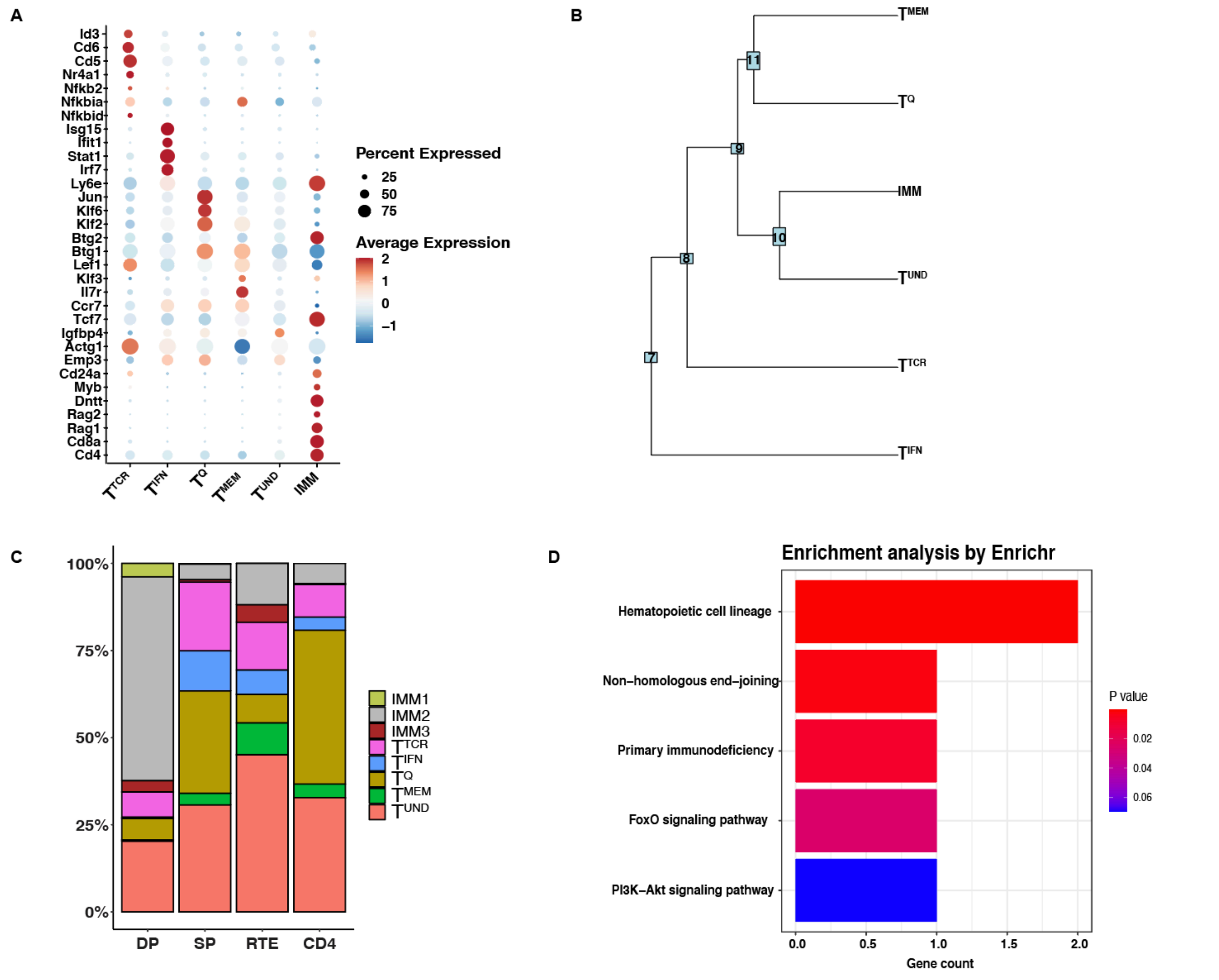


**Figure S9. Thymic derived naïve CD4$^+$ T cell clusters are maintained in recent thymic emigrants in the periphery. A:** Dot plot showing thymic naïve DP, SP and peripheral splenic CD4$^+$ T cell markers, including the markers for the thymic immature cluster. **B:** Cluster dendrogram showing similarities between clusters. **C:** Frequency plot of thymic DP and thymic SP, peripheral splenic naïve CD4$^+$ T cells and peripheral RAG2-GFP$^+$ naïve CD4$^+$ T cells that represented Recent Thymic Emigrants (RTEs). RTEs were enriched using RAG2-GFP mice. **D:** EnrichR plot using DEGs in the Immature cluster of naïve DP, SP and peripheral naïve CD4$^+$ T cells. Mice used for DP (thymic), SP (thymic) and CD4 (splenic) groups were VISTA CD4-Cre-, representative of wild type mice. RTE refers to GFP$^+$ splenic naïve CD4$^+$ T cells from RAG2-GFP mice. (DP n=1, SP n=2, RTE n=2, CD4 n=3).

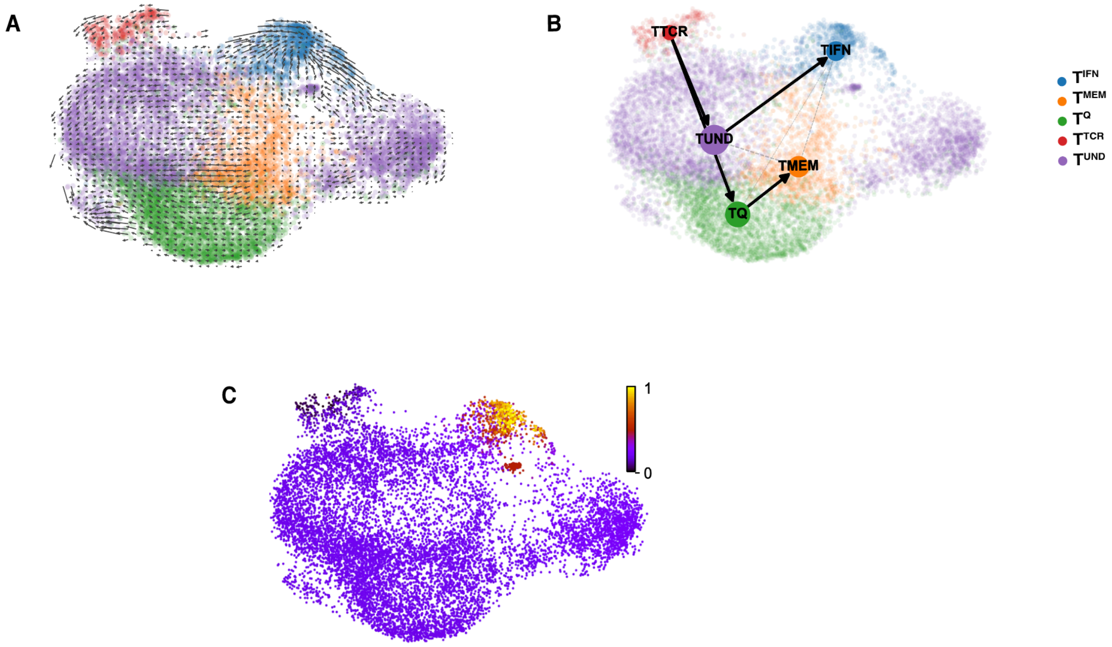


**Figure S10. Pseudotime shows stability and plasticity of naïve CD4$^+$ T cell clusters. A:** RNA velocity of splenic naïve CD4+ T cells. **B:** PAGA plot from RNA velocity analysis of splenic naïve CD4+ T cells. **C:** Pseudotime plot showing the lineage trajectory of splenic naïve CD4$^+$ T cells.

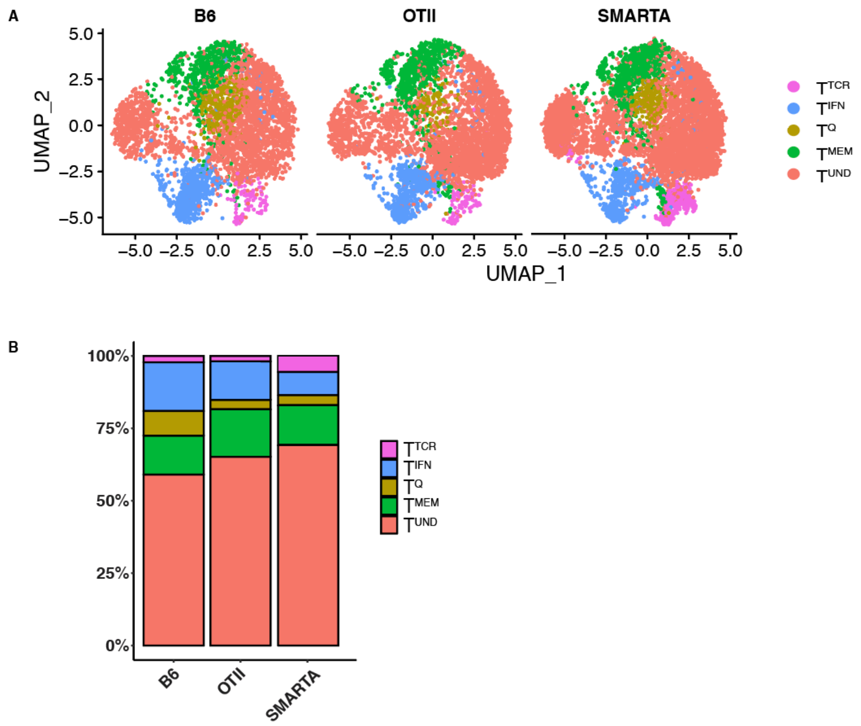


**Figure S11. Naïve CD4⁺ T cell clusters from monoclonal vs polyclonal T cells are not transcriptionally distinct. A:** UMAPs showing naïve CD4⁺ transcriptional clusters from TCR transgenics OT-II and SMARTA and control B6 mice. (n=2). **B:** Frequency plots quantifying clusters from TCR transgenics OT-II and SMARTA and control B6 mice.

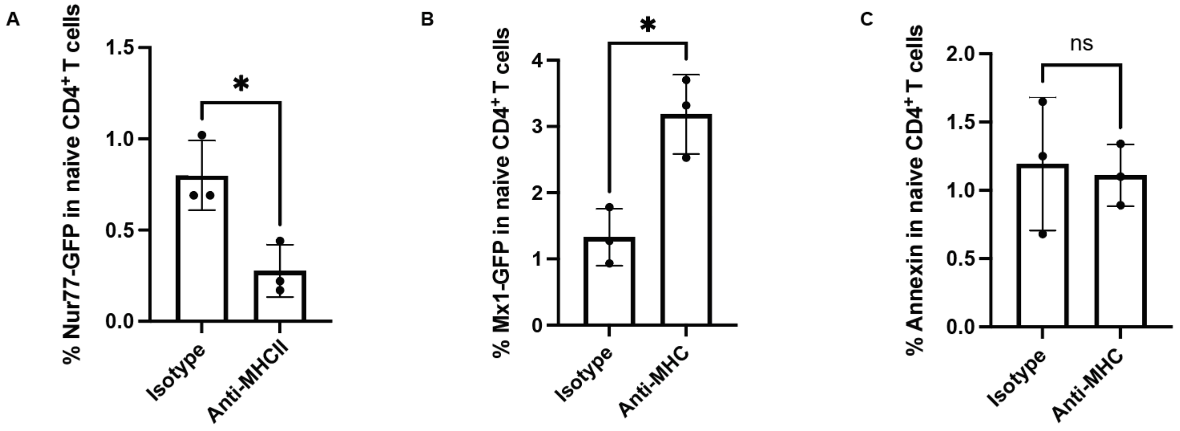


**Figure S12. Naïve CD4+ T cells from MHCII or control isotype treated mice. A.** Nur77-GFP, **B.** Mx1-GFP and **C.** Annexin staining in naïve CD4+ T cells treated with anti-MHCII or isotype control. Significance testing performed using unpaired t-test. * <0.05, ** <0.01, ***<0.001. (n=3).

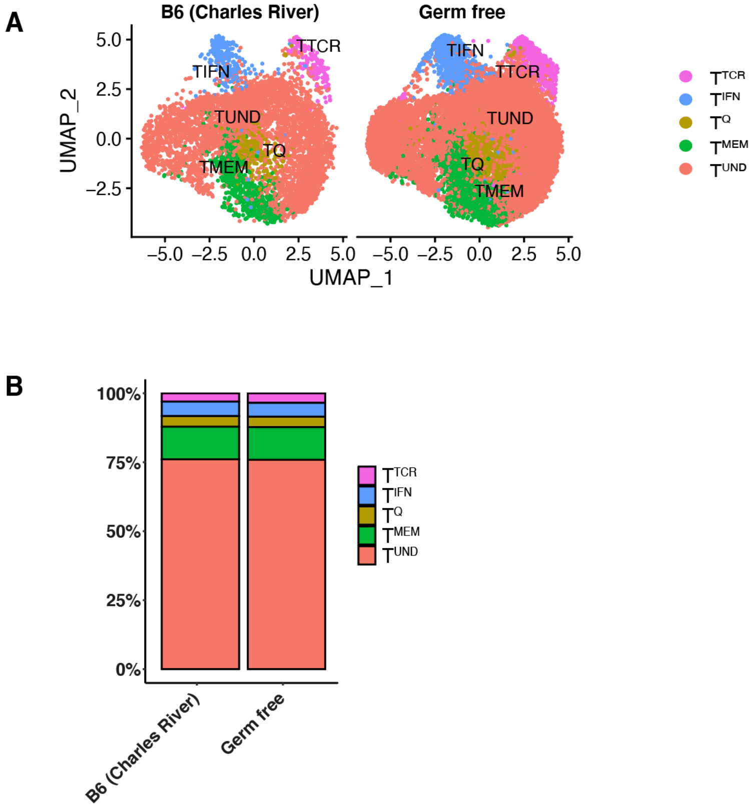


**Figure S13. Microbiome experience does not influence naïve CD4⁺ T cell transcriptional heterogeneity. A:** UMAP showing clustering of naïve CD4⁺ T cells from germ free and specific pathogen free mice. (n=2). **B:** Frequency plot qualifying the UMAP frequencies in (A).

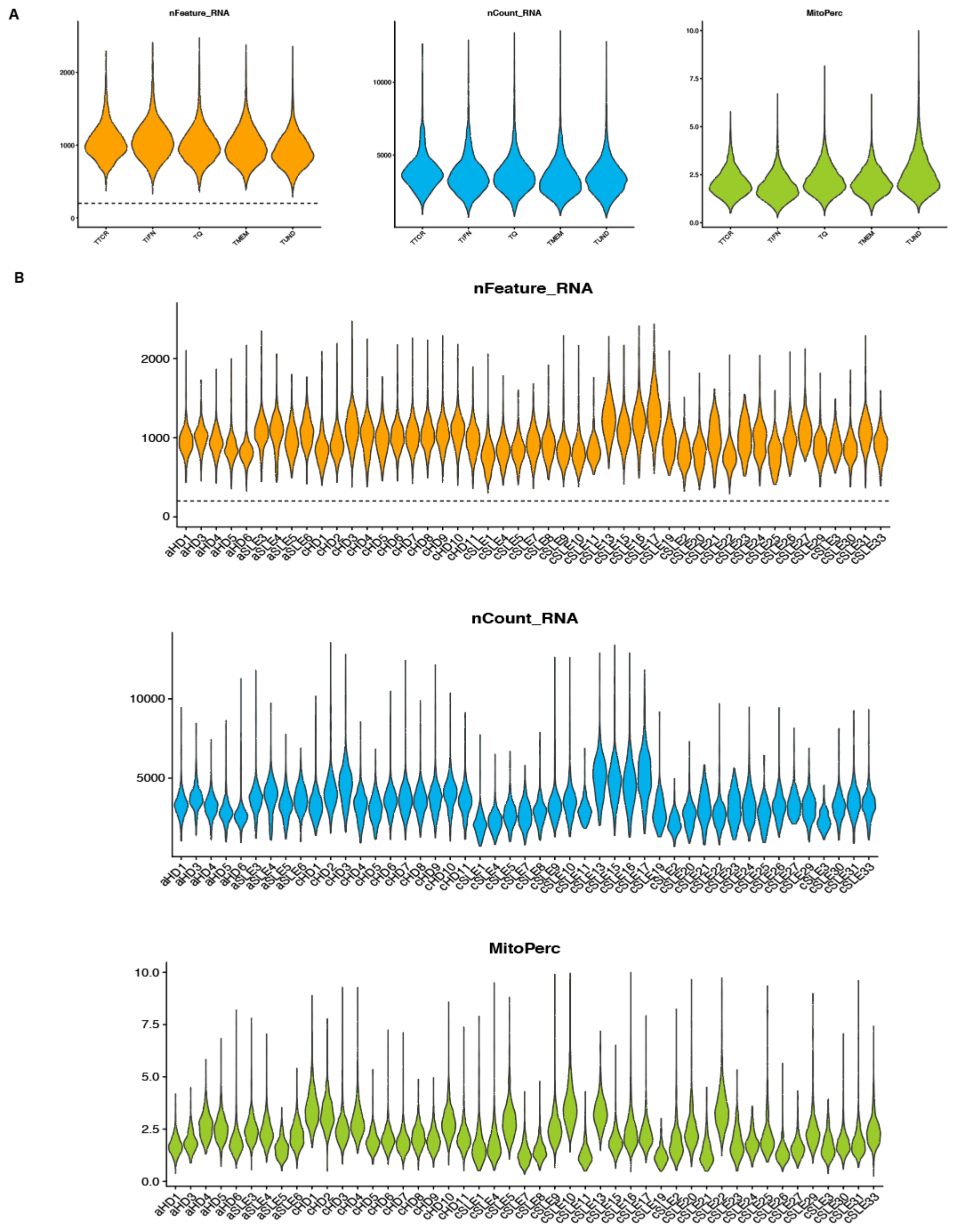


**Figure S14. Representative plots for human single cell RNA sequencing quality control pipeline showing nFeature, nCount and mitochondrial content. A**. nFeature, nCount and mitochondrial content per cluster after quality control filtering. **B.** nFeature, nCount and mitochondrial content separated out donor.

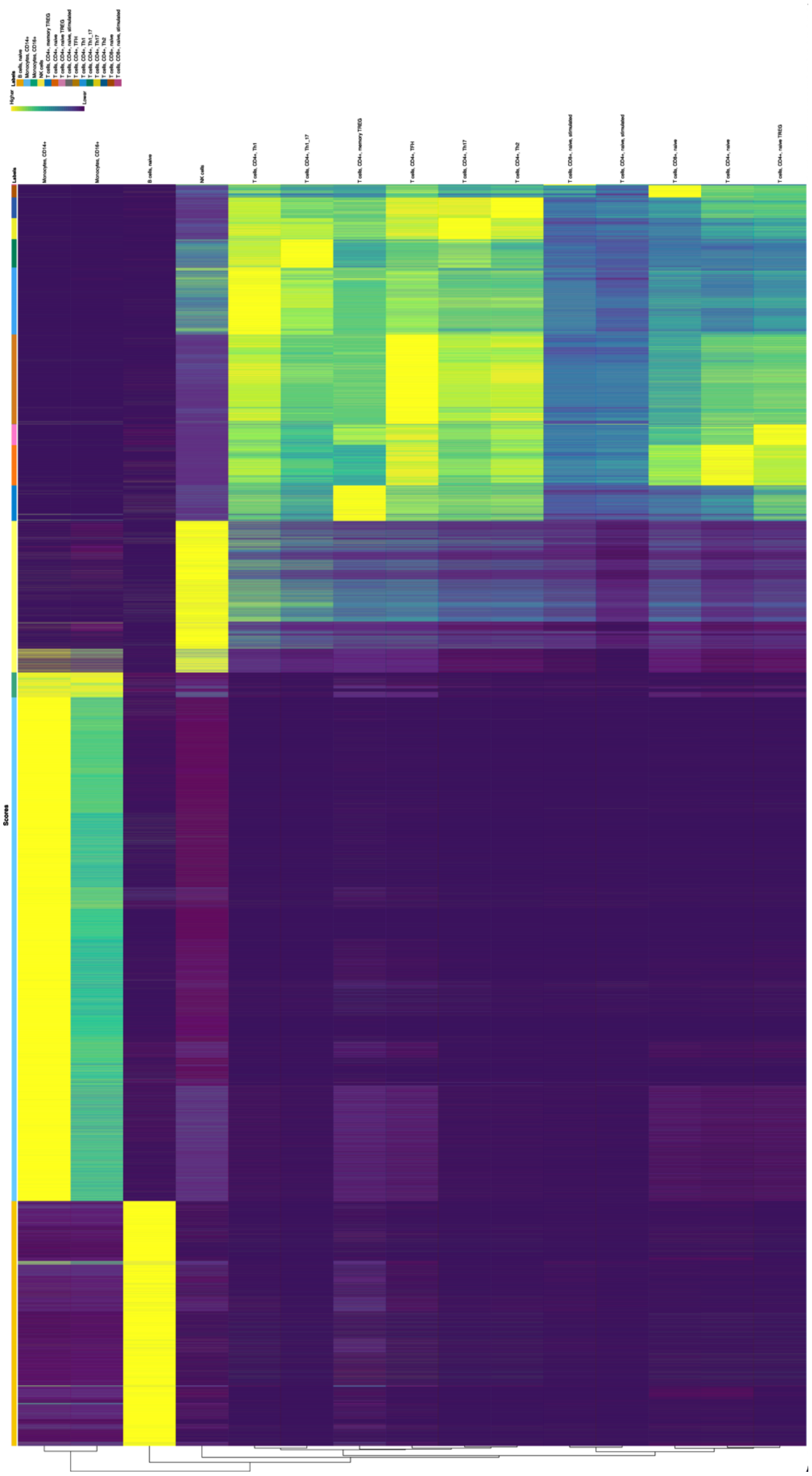

**Figure S15. SingleR predictions for immune cell subsets in Tabula Sapiens dataset.**

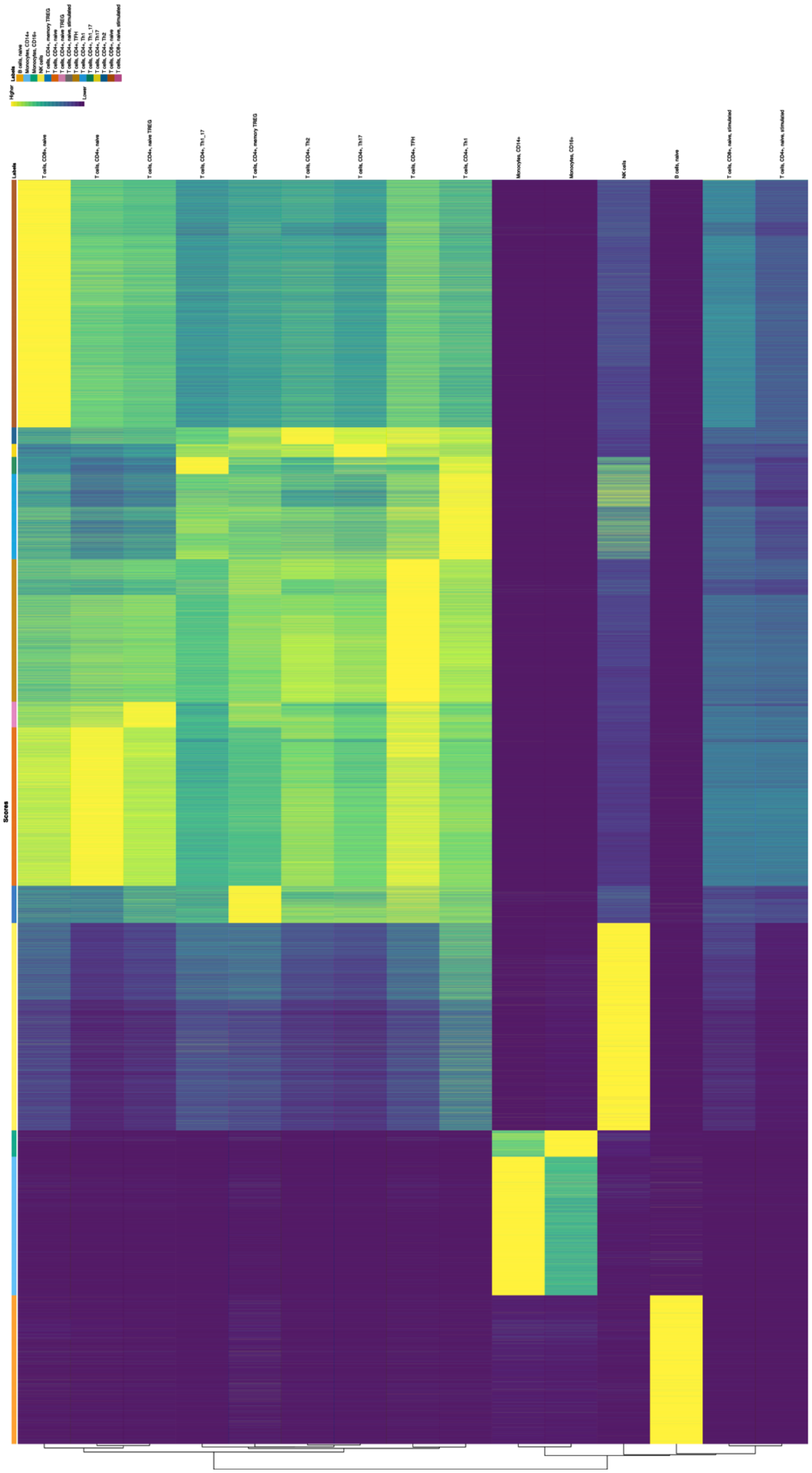

**Figure S16. SingleR predictions for immune cell subsets in lupus dataset.**

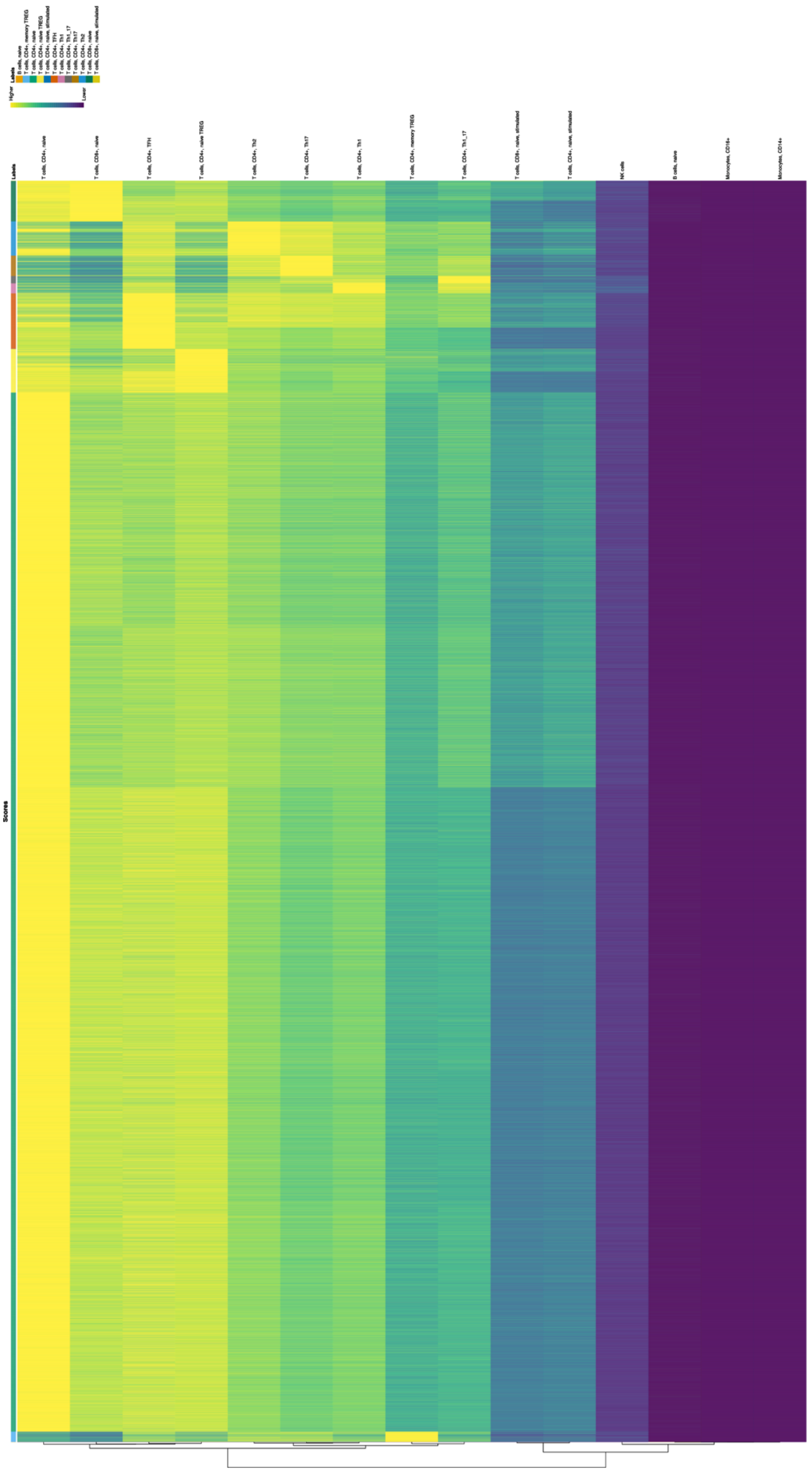


**Figure S17. SingleR predictions for immune cell subsets in DHMC in-house sorted naïve CD4⁺ T cells.**

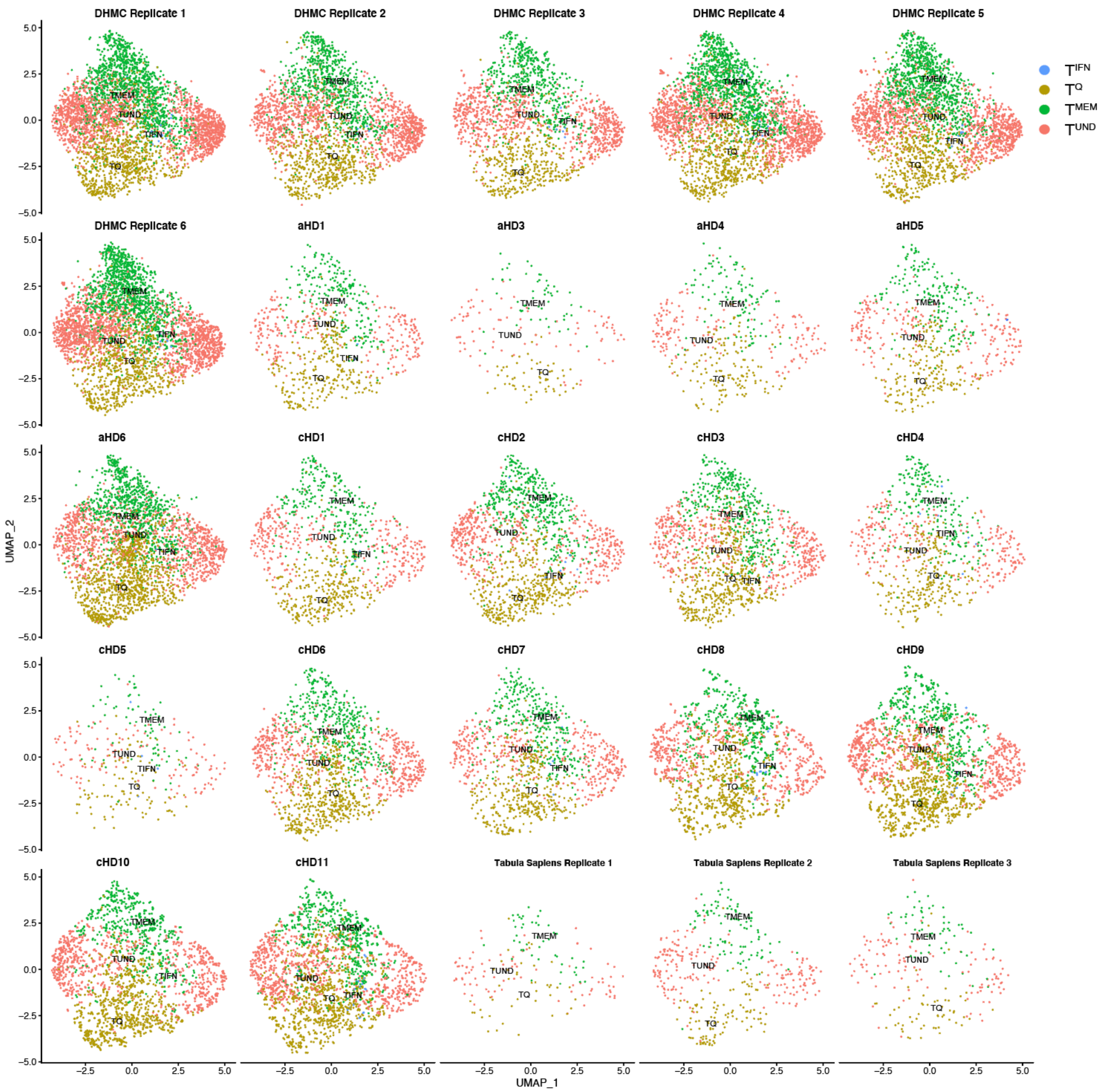


**Figure S18. UMAPs separated by donor for human healthy dataset (DHMC, lupus and tabula sapiens datasets).**

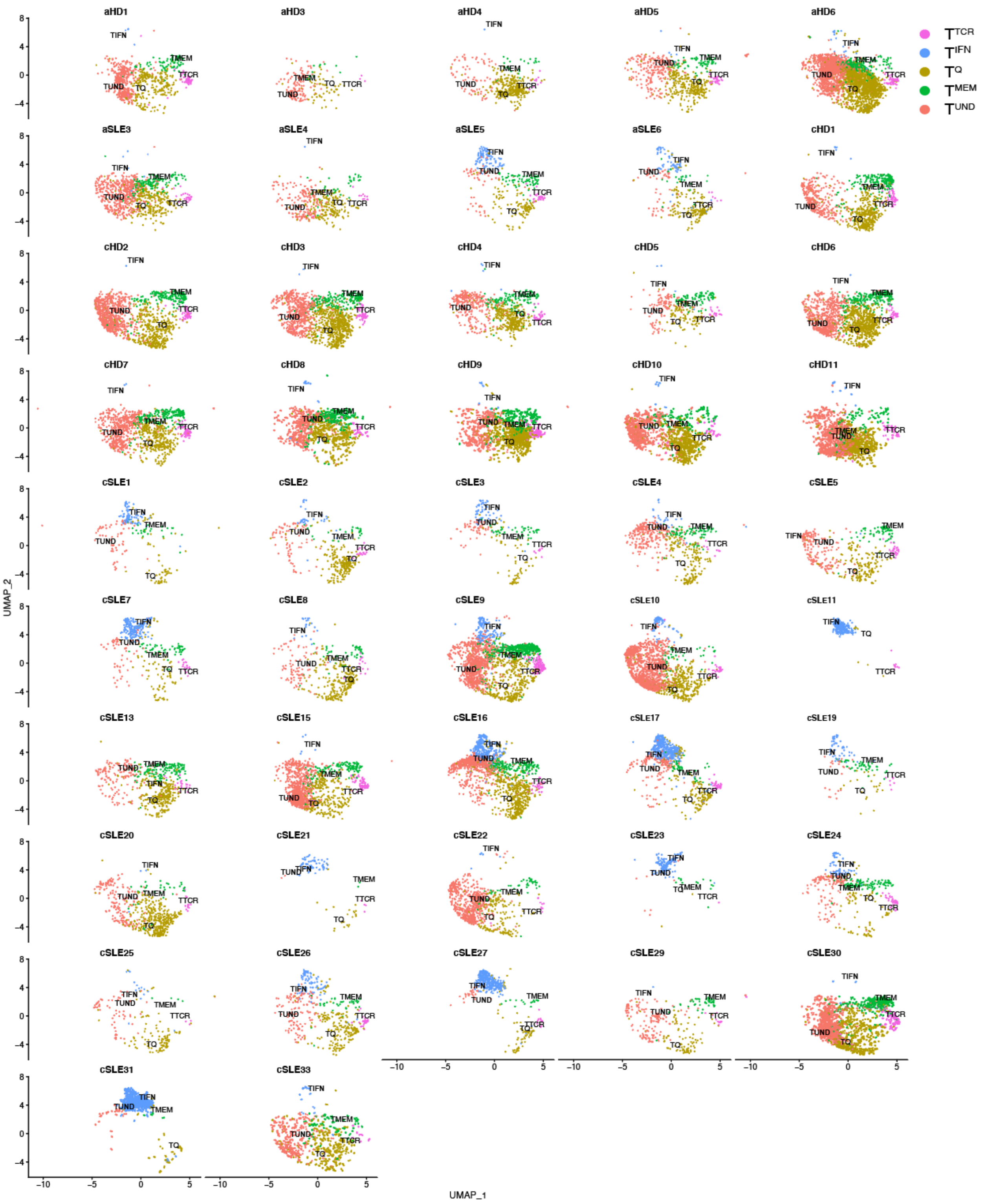

Figure S19. UMAPs separated by donor for lupus dataset.

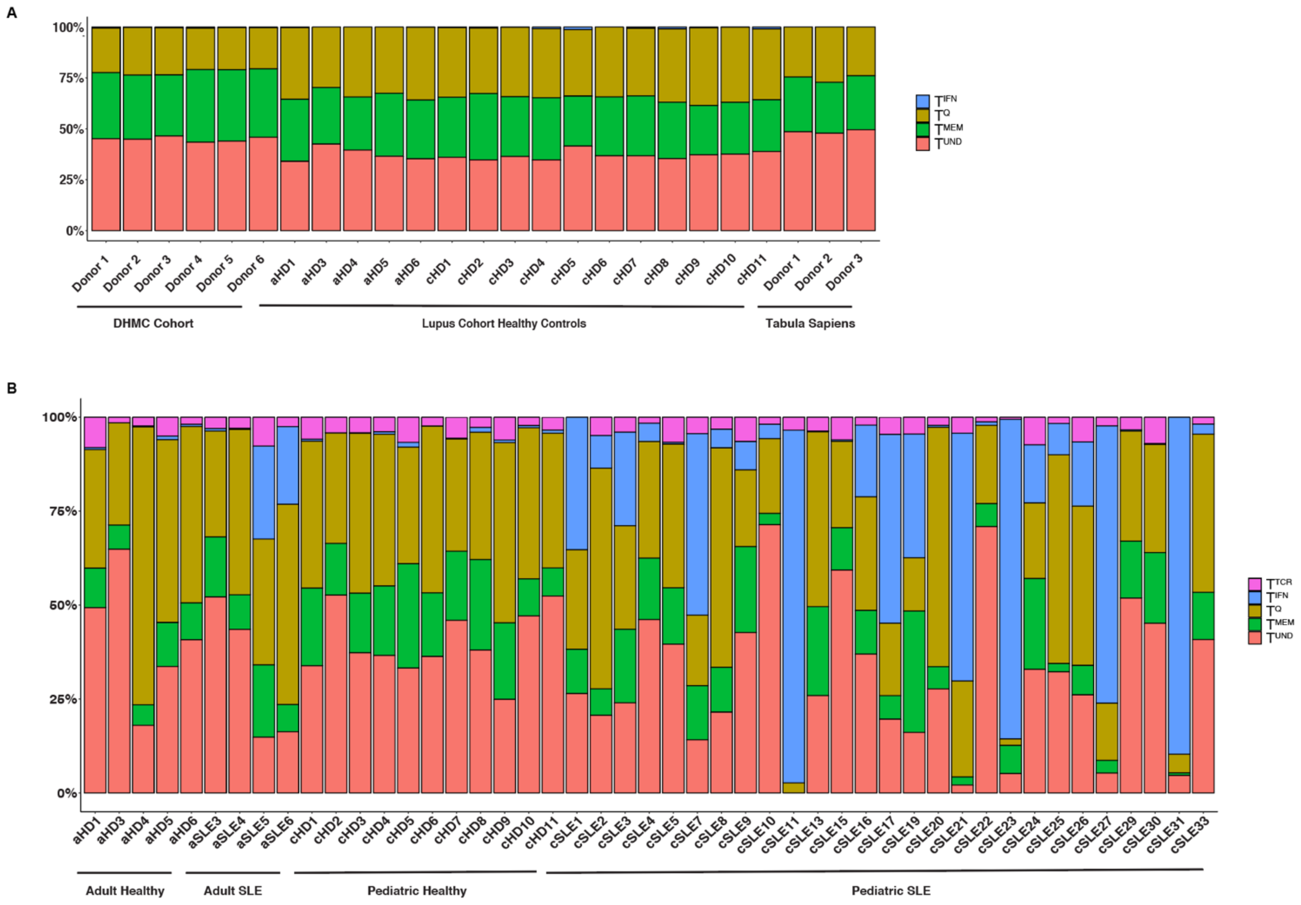


**Figure S20. Variations in naïve $CD4^+$ T cell clusters across donors in healthy and lupus datasets shows loss in stability in naïve $CD4^+$ T cell clusters in lupus. A:** Naïve $CD4^+$ T cell transcriptional clusters by donor in 3 healthy datasets - our DHMC cohort, healthy controls from the lupus dataset and Tabula Sapiens dataset. **B:** Naïve $CD4^+$ T cell transcriptional clusters by donor in the lupus datasets, showing both healthy and lupus experiencing pediatric and adult patients. Donors with naïve $CD4^+$ T cell numbers less than 100 were discarded from analysis.

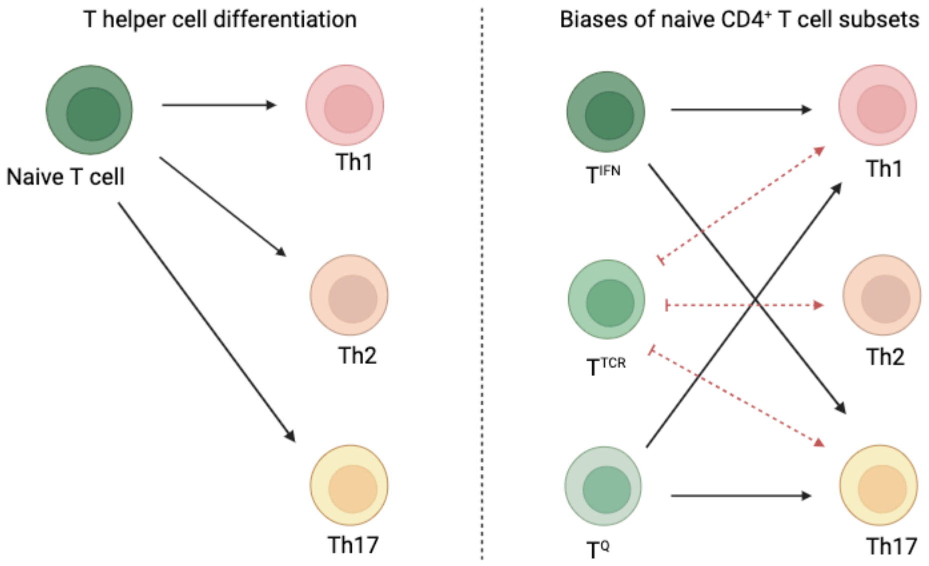


**Figure S21. Model depicting the redefined lineage biases dependent on naïve $CD4^+$ T cell cluster identity.** Black arrows represent skewing biases and red dotted arrows represent inhibition of differentiation trajectories. Figure created in BioRender.

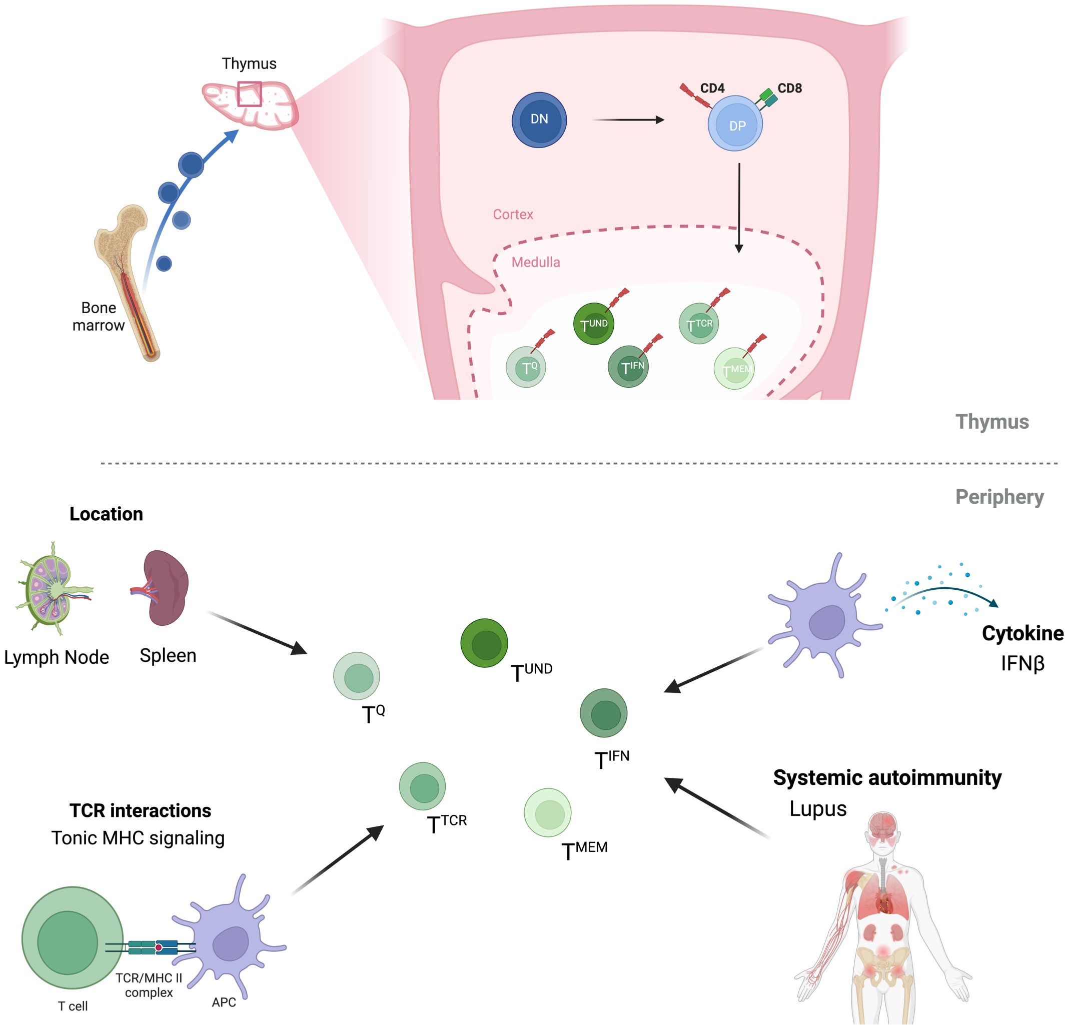


**Figure S22. Summary of the environmental cues that impact naïve CD4$^+$ T cell subsets.** The thymus impacts the emerge of all clusters during T cell development. In the periphery, tonic signaling controls $T^{TCR}$, tissue microenvironment impacts $T^{Q}$, while lupus and IFNβ influence $T^{IFN}$. Figure created in BioRender.

| | avg_log2FC | p_val_adj | cluster | gene |
|---|---|---|---|---|
| **Isg15.2** | 1.954369158 | 0 | TIFN | Isg15 |
| **Ifi27l2a.2** | 1.650667835 | 1.52E-205 | TIFN | Ifi27l2a |
| **Bst2.3** | 1.588458876 | 2.47E-224 | TIFN | Bst2 |
| **Irf7.3** | 1.526434888 | 1.31E-240 | TIFN | Irf7 |
| **Ly6a.3** | 1.50364345 | 3.28E-186 | TIFN | Ly6a |
| **Ifit3** | 1.250361205 | 0 | TIFN | Ifit3 |
| **Stat1.3** | 1.206833125 | 2.44E-158 | TIFN | Stat1 |
| **Zbp1.3** | 1.005371713 | 4.72E-176 | TIFN | Zbp1 |
| **Slfn5** | 0.980556081 | 1.98E-167 | TIFN | Slfn5 |
| **Isg20** | 0.938249423 | 2.87E-181 | TIFN | Isg20 |
| **Ifit1** | 0.916530793 | 1.64E-249 | TIFN | Ifit1 |
| **Ly6c1.1** | 0.91420587 | 1.28E-52 | TIFN | Ly6c1 |
| **Slfn1.1** | 0.889972139 | 4.56E-102 | TIFN | Slfn1 |
| **Rnf213.1** | 0.853788163 | 6.93E-136 | TIFN | Rnf213 |
| **Ifi47.1** | 0.844785083 | 1.21E-85 | TIFN | Ifi47 |
| **Ly6e.1** | 0.842306812 | 5.11E-95 | TIFN | Ly6e |
| **Ifi203.1** | 0.802468333 | 1.94E-87 | TIFN | Ifi203 |
| **H2-T23** | 0.777907907 | 2.33E-87 | TIFN | H2-T23 |
| **Rtp4** | 0.777299303 | 9.01E-148 | TIFN | Rtp4 |
| **Daxx** | 0.771964491 | 1.58E-107 | TIFN | Daxx |
| **Usp18** | 0.763124998 | 9.28E-177 | TIFN | Usp18 |
| **Samhd1.1** | 0.695226717 | 2.54E-52 | TIFN | Samhd1 |
| **Ifi213** | 0.684699729 | 1.69E-93 | TIFN | Ifi213 |
| **Phf11b** | 0.676366775 | 2.99E-98 | TIFN | Phf11b |
| **Xaf1** | 0.67295841 | 1.69E-115 | TIFN | Xaf1 |
| **H2-T22** | 0.661115955 | 4.59E-63 | TIFN | H2-T22 |
| **Ms4a4b.2** | 0.619795816 | 1.76E-51 | TIFN | Ms4a4b |
| **AW112010** | 0.607360432 | 8.41E-34 | TIFN | AW112010 |
| **Gbp6** | 0.600023679 | 1.69E-136 | TIFN | Gbp6 |
| **Oasl2** | 0.596644306 | 9.56E-154 | TIFN | Oasl2 |
| **Mndal** | 0.589671837 | 9.94E-52 | TIFN | Mndal |
| **Igtp** | 0.577020786 | 3.79E-74 | TIFN | Igtp |
| **Sp100** | 0.559817726 | 1.9E-42 | TIFN | Sp100 |
| **Tapbp** | 0.559738193 | 1.36E-49 | TIFN | Tapbp |
| **Trim30a** | 0.554844423 | 8.12E-77 | TIFN | Trim30a |
| **Irgm1** | 0.542305958 | 5.66E-85 | TIFN | Irgm1 |
| **Epsti1** | 0.535521203 | 4.29E-49 | TIFN | Epsti1 |
| **Psmb8.1** | 0.530759273 | 2.75E-44 | TIFN | Psmb8 |
| **Psme1** | 0.515398826 | 4.11E-47 | TIFN | Psme1 |
| **Shisa5** | 0.514877376 | 4.43E-60 | TIFN | Shisa5 |
| **Lgals9** | 0.510713142 | 2.2E-39 | TIFN | Lgals9 |
| **Ifi208** | 0.508911793 | 5.06E-53 | TIFN | Ifi208 |
| **Il7r** | 0.665623842 | 8.08E-53 | TMEM | Il7r |
| **Crlf3** | 0.561841005 | 3.22E-54 | TMEM | Crlf3 |
| **Cnn2** | -0.454840783 | 5.46E-29 | TMEM | Cnn2 |
| **Pfn1** | -0.568768864 | 9.63E-78 | TMEM | Pfn1 |
| **Cd52** | -0.60276123 | 6.36E-63 | TMEM | Cd52 |
| **Actg1** | -0.684903763 | 4.79E-73 | TMEM | Actg1 |
| **Gm42418** | -0.693510769 | 9.29E-05 | TMEM | Gm42418 |
| **Actb** | -0.705063757 | 5.86E-142 | TMEM | Actb |
| **Jun.2** | 0.892844921 | 9.28E-103 | TQ | Jun |
| **Bst2.2** | -0.48556722 | 7.24E-14 | TQ | Bst2 |
| **Isg15.1** | -0.626217104 | 1.18E-18 | TQ | Isg15 |
| **Ifi27l2a.1** | -0.646707547 | 3.66E-23 | TQ | Ifi27l2a |
| **Gm42418.2** | -0.807795476 | 9.41E-15 | TQ | Gm42418 |
| **Nr4a1** | 1.381368859 | 5.31E-33 | TTCR | Nr4a1 |
| **Nfkbia.2** | 1.10989336 | 1.41E-08 | TTCR | Nfkbia |
| **Cd5** | 0.964813922 | 1.61E-15 | TTCR | Cd5 |
| **Itm2a** | 0.923743811 | 7.92E-06 | TTCR | Itm2a |
| **Nfkbid** | 0.909659257 | 2.1E-28 | TTCR | Nfkbid |
| **Egr1** | 0.852550171 | 7.65E-10 | TTCR | Egr1 |
| **Eif5a** | 0.846708958 | 1.91E-12 | TTCR | Eif5a |
| **Id3** | 0.816114947 | 1.87E-10 | TTCR | Id3 |
| **Ptma** | 0.707212947 | 6.03E-13 | TTCR | Ptma |
| **Srgn.1** | 0.660628171 | 0.012093841 | TTCR | Srgn |
| **Eif4a1** | 0.654819458 | 4.06E-07 | TTCR | Eif4a1 |
| **Cd69.1** | 0.65216419 | 0.066856179 | TTCR | Cd69 |
| **Ptpn6.1** | 0.629952363 | 1E-11 | TTCR | Ptpn6 |
| **Hsp90ab1** | 0.61920352 | 9.7E-11 | TTCR | Hsp90ab1 |
| **Ncl** | 0.540566264 | 1 | TTCR | Ncl |
| **Cd6** | 0.540028559 | 2.7E-08 | TTCR | Cd6 |
| **Ppp1r14b** | 0.535649395 | 2.67E-07 | TTCR | Ppp1r14b |
| **Myc** | 0.529589526 | 1 | TTCR | Myc |
| **Foxp4** | 0.527072885 | 3.59E-12 | TTCR | Foxp4 |
| **Ldha** | 0.526681471 | 5.76E-05 | TTCR | Ldha |
| **Dusp2** | 0.525934497 | 0.009528252 | TTCR | Dusp2 |
| **Npm1** | 0.522404946 | 0.195582211 | TTCR | Npm1 |
| **Zfp36** | 0.519290129 | 1 | TTCR | Zfp36 |
| **Il2rg** | 0.506001938 | 0.002012912 | TTCR | Il2rg |
| **Cd82** | 0.501199636 | 1 | TTCR | Cd82 |
| **Nme1** | 0.500127736 | 0.007269548 | TTCR | Nme1 |
| **Slfn1.2** | -0.463631773 | 0.327414274 | TTCR | Slfn1 |
| **Rflnb.1** | -0.465624184 | 0.020700893 | TTCR | Rflnb |
| **Ighm** | -0.473082704 | 0.03214052 | TTCR | Ighm |
| **Igfbp4.3** | -0.603243782 | 0.464699796 | TTCR | Igfbp4 |
| **Klf2.2** | -0.641160193 | 0.000214479 | TTCR | Klf2 |
| **Tsc22d3.1** | -0.662586831 | 9.26E-06 | TTCR | Tsc22d3 |
| **Txnip** | -0.811289022 | 8.08E-07 | TTCR | Txnip |
| **Gm42418.1** | 0.823264781 | 3.3E-29 | TUNDIFF | Gm42418 |
| **Jun.1** | -0.532774216 | 9.79E-41 | TUNDIFF | Jun |
| **Ly6a.1** | -0.567234612 | 1.24E-26 | TUNDIFF | Ly6a |
| **Irf7.1** | -0.569221822 | 1.39E-33 | TUNDIFF | Irf7 |
| **Bst2.1** | -0.575451906 | 7.52E-22 | TUNDIFF | Bst2 |
| **Ifi27l2a** | -0.596935882 | 3.87E-22 | TUNDIFF | Ifi27l2a |
| **Isg15** | -0.742313603 | 1.98E-35 | TUNDIFF | Isg15 |

**Table S1: Markers for mouse single cell clusters.**

| | avg_log2FC | p_val_adj | cluster | gene |
|---|---|---|---|---|
| **IFI6** | 1.155015875 | 4.9E-45 | TIFN | IFI6 |
| **ISG15** | 1.079970094 | 2.48E-35 | TIFN | ISG15 |
| **STAT1** | 1.049918715 | 2.35E-37 | TIFN | STAT1 |
| **IFITM1.1** | 0.866528588 | 3.39E-07 | TIFN | IFITM1 |
| **MX1** | 0.792242784 | 2.59E-42 | TIFN | MX1 |
| **IFI44L** | 0.75779067 | 8.05E-118 | TIFN | IFI44L |
| **LY6E** | 0.752273832 | 7.78E-12 | TIFN | LY6E |
| **MT2A** | 0.711845438 | 5.72E-16 | TIFN | MT2A |
| **XAF1** | 0.703793409 | 6.76E-33 | TIFN | XAF1 |
| **TRIM22** | 0.632261399 | 8.65E-12 | TIFN | TRIM22 |
| **UBE2L6** | 0.506859548 | 3.23E-13 | TIFN | UBE2L6 |
| **FOS.1** | -0.575702234 | 2.92E-191 | TMEM | FOS |
| **JUNB.1** | -0.879717655 | 0 | TMEM | JUNB |
| **JUNB.2** | 1.249266644 | 0 | TQ | JUNB |
| **FOS.2** | 0.931396173 | 0 | TQ | FOS |
| **JUN.2** | 0.919902759 | 0 | TQ | JUN |
| **DUSP1.2** | 0.693638663 | 0 | TQ | DUSP1 |
| **NFKBIA.2** | 0.559120839 | 0 | TQ | NFKBIA |
| **CD69.1** | 0.553869139 | 0 | TQ | CD69 |
| **CXCR4.1** | 0.543796782 | 0 | TQ | CXCR4 |
| **ZFP36.1** | 0.503991548 | 0 | TQ | ZFP36 |
| **MTRNR2L12.1** | -0.500473614 | 4.33E-154 | TQ | MTRNR2L12 |
| **JUNB** | -0.563059958 | 7.32E-193 | TUND | JUNB |

**Table S2: Markers for human single cell clusters.**

| | | Mouse | | | | |
|---|---|---|---|---|---|---|
| | | TIFN | TMEM | TQ | TTCR | TUND |
| Human | TIFN | 0.35615371 | -0.2257231 | -0.24556111 | 0.06411807 | -0.21393918 |
| | TMEM | 0.04725379 | 0.4243488 | 0.06068659 | -0.32741842 | 0.11802961 |
| | TQ | -0.2650143 | 0.0552417 | 0.28617781 | 0.04471883 | -0.06048239 |
| | TUND | -0.25504152 | -0.1611469 | -0.0390069 | 0.18210937 | 0.2643629 |

**Table S3: R values for Mouse-Human naïve $CD4^+$ T cell cluster correlation.**
